# Understanding the Hidden Complexity of Latin American Population Isolates

**DOI:** 10.1101/340158

**Authors:** Jazlyn A. Mooney, Christian D. Huber, Susan Service, Jae Hoon Sul, Clare D. Marsden, Zhongyang Zhang, Chiara Sabatti, Andrés Ruiz-Linares, Gabriel Bedoya, Nelson Freimer, Costa Rica/Colombia Consortium for Genetic Investigation of Bipolar Endophenotypes, Kirk E. Lohmueller

## Abstract

Most population isolates examined to date were founded from a single ancestral population. Consequently, there is limited knowledge about the demographic history of admixed population isolates. Here we investigate genomic diversity of recently admixed population isolates from Costa Rica and Colombia and compare their diversity to a benchmark population isolate, the Finnish. These Latin American isolates originated during the 16^th^ century from admixture between a few hundred European males and Amerindian females, with a limited contribution from African founders. We examine whole genome sequence data from 449 individuals, ascertained as families to build mutigenerational pedigrees, with a mean sequencing depth of coverage of approximately 24X. We find that Latin American isolates have increased genetic diversity relative to the Finnish. However, there is an increase in the amount of identity by descent (IBD) segments in the Latin American isolates relative to the Finnish. The increase in IBD segments is likely a consequence of a very recent and severe population bottleneck during the founding of the admixed population isolates. Furthermore, the proportion of the genome that falls within a long run of homozygosity (ROH) in Costa Rican and Colombian individuals was significantly greater than that in the Finnish, suggesting more recent consanguinity in the Latin American isolates relative to that seen in the Finnish. Lastly, we found that recent consanguinity increased the number of deleterious variants found in the homozygous state, which is relevant if deleterious variants are recessive. Our study suggests there is no single genetic signature of a population isolate.

## Introduction

The use of population isolates to map Mendelian and complex diseases has been a key feature of medical genomics. In addition to experiencing the bottleneck involved with the migration out of Africa, some populations underwent subsequent bottlenecks and remained in relative seclusion afterward. These populations formed present-day isolates^1^. There are numerous benefits to use population isolates to map genes underlying disease. First, and perhaps the largest benefit, is the increased homogeneity of genomes in isolates when compared to outbred populations. Second, isolates experience greater genetic drift than the population from which they were founded. Drift can allow disease causing alleles to exist at an appreciable frequency in isolated populations. Third, isolates may be endogamous and the cryptic relatedness of individuals leads to an enrichment in prevalence of the phenotype of interest and an enrichment of homozygous disease variants^1–4^. Lastly, isolates may also have increased cultural and environmental homogeneity, resulting in a reduction of variability in phenotype due to non-genetic sources.

Generally, the genomes of population isolates are thought to exhibit several hallmark features of genetic diversity. Due to bottlenecks associated with their founding, it is thought that isolates should carry lower levels of genetic diversity and lower haplotype diversity than closely related non-isolated populations. Drift experienced by isolates is magnified by small population size, which generates more linkage disequilibrium (LD) than in non-isolated populations. In addition to increased LD, individuals from isolated populations tend to share more regions of the genome identical by descent (IBD) due to small population sizes. Further, due to the isolation after founding and recent mating practices, isolates may have larger regions of the genome found in runs of homozygosity (ROHs) as a result of recent inbreeding. Lastly, bottlenecks and inbreeding should impact patterns of deleterious variation^5–7^. Consequently, one would predict that individuals from isolates will have fewer segregating sites, and the remaining deleterious variants will be segregating at a higher frequency^8^. Indeed, genomic studies over the last decade have documented several of these signatures^2,9,10^. However, it is known that not all isolates share the same demographic history. Therefore, it is essential that we understand how the factors shaping genetic variation in a population, are influenced by the unique demographic history of the population.

One archetypal human population isolate with a demography that has been extensively studied is the Finnish ^2,11–13^. Finland was populated through two separate major migrations. The first wave originated 4000 years ago from the west, and the second wave originated from the southern shores approximately 2000 years ago. There was a subsequent internal migration and expansion around the 16^th^ century. Briefly, the small number of founders, relative isolation, serial bottlenecks, and recent expansion in Finland has allowed drift to play a large role in shaping the gene pool of this population. The aforementioned demographic history of Finland has led to an increase in the prevalence of rare heritable Mendelian diseases, which has made this population particularly fruitful for disease gene discovery studies^12,14^. Most of the initial disease gene discovery studies in Finland exploited LD mapping in affected families and well curated genealogical records to identify causal and candidate variants^12^. More recently, it has been possible to apply population-based linkage analyses to identify disease associated variants as an alternative to GWAS (unpublished data)^15^ due to the availability of whole genome sequence data in conjunction with extensive electronic health records.

A number of studies have shown that disease detection power can be improved by studying population isolates other than the Finnish^10,16–18^. For example, the Greenlandic Inuit (GI) experienced an extreme bottleneck which caused a depletion of rare variants and segregating sites in their genome^18^. The remaining segregating variants are maintained at higher allele frequencies and a larger proportion of these SNPs are deleterious when compared to non-isolated populations. Another study on Southeast Asian populations showed similar results. Specifically, South East Asian populations have experienced more severe founder effects than the Finnish^17^, thus causing an excess of rare alleles associated with recessive disease. A study of European population isolates compared the isolates with the closest non-isolated population from similar geographic regions^10^, and found that the total number of segregating sites was depleted across all isolates relative to the comparison non-isolate. Of the sites that were segregating in isolates, between ~30,000-122,000 sites existed at an appreciable frequency (MAF > 5.6%), while remaining rare (MAF < 1.4%) in all of the non-isolate population samples. These variants could serve as candidate markers in genome-wide association studies (GWAS) for novel associations and included SNPs that had been previously associated with cardio-metabolic traits.

As previously mentioned, population isolates tend to have a depletion of segregating sites and an increased number of homozygous sites in their genomes. Pemberton and colleagues^19,20^ demonstrated that levels of homozygosity differ across the globe, and that long ROH, 2Mb or greater, are strongly influenced by very recent demography. A study involving multiple Jewish isolates showed a link between historic consanguinity and the amount of long ROH in the genome^21^. The same correlation between long ROH and parental consanguinity was also observed in Middle Eastern isolates^4^. In another study, authors observed an increased count of ROH and total length of the genome within an ROH in Greek isolates from the Pomak villages and the island of Crete relative to a non-isolated Greek cohort^16^. Lastly, a study in Sardinians showed there was sub-structure within the island when comparing the amount of long ROH across sampled populations^22^. Authors identified an enrichment of long ROH in sub-populations that had experienced recent endogamy, while sub-populations that had not experienced such isolation have ROH levels similar to that of non-Sardinian Italians.

While there have been many studies of genetic variation in population isolates, the studies described above have focused on populations where the founders all came from the same ancestral population. However, the founders of Latin American population isolates have come from distinct continental populations. We sampled individuals from mountainous regions of Costa Rica and Colombia where geographic barriers resulted in populations remaining isolated since their founding in the 16^th^ and 17^th^ centuries, until the mid-20^th^ century^23^. Both groups share a similar demographic history, having originated primarily from admixture between a few hundred European males and Amerindian females, with a limited contribution from African founders. After the founding event, both populations experienced a subsequent bottleneck and then a recent expansion, within the last 300 years, the expansion increased the population size over 1000-fold since the initial founding event^23^. The effect that admixture has had on overall patterns of genetic variation in isolates remains elusive, and it is unclear whether these populations share the typical genomic signatures seen in population isolates. While the small founding population size could reduce diversity, because the Costa Rican and Colombian isolates were founded from multiple diverse populations, they could potentially have increased in diversity relative to other population isolates. Lastly, the impact of admixture on deleterious variation remains unclear.

To better understand patterns of genetic variation in admixed isolated populations, we compared the Colombian and Costa Rican population isolates to a benchmark isolate, the Finnish, as well as other 1000 Genomes Project populations^24^. We observe that relative to the Finnish, Latin American isolates have increased genetic diversity but an excess of IBD segments. Moreover, we detect an increase in the proportion of an individual’s genome that falls within a long ROH in Latin American isolates relative to all other sampled populations and an enrichment of deleterious variation within these long ROH. Demographic simulations indicate that the enrichment of long ROH is a consequence of recent inbreeding in Latin American isolates. We corroborate these results by leveraging extended pedigree data. Pedigree inbreeding coefficients explain approximately 21.8% of the amount of an individual’s genome within a long ROH. Next, we examine the relationship between the proportion of European, Native American, and African ancestry and the amount of the genome within an ROH, as well as the relationship to an individual’s pedigree inbreeding coefficient. To our knowledge, this is the first time long ROH have been examined in admixed isolated populations. Further, we examine demography across both recent and ancient timescales in these isolates. Our work sheds light on how the distinct demographic histories of population isolates affect both genetic diversity and the distribution of deleterious variation across the genome.

## Methods

### Pedigree Data for Costa Rican and Colombian Individuals

Our study included 10 Costa Rican (CR) and 12 Colombian (CO) multi-generational pedigrees ascertained to include individuals affected by Bipolar Disorder 1. More extensive details about the curation of pedigree data and clinical assessments of diagnosis can be found in Fears et al.^25^.

### Identifying Unrelated Individuals

We defined unrelated individuals as those who are at most third-degree relatives. We chose this threshold of relatedness because the families from CR and CO are known to be cryptically related. We used KING^26^ to identify 30 unrelated individuals from CR and CO. 24 of the 30 unrelated individuals in the CO are founders in the pedigree and 15 of the 30 unrelated individuals in the CR are founders. The algorithm implemented in KING estimates familial relationships by modeling the genetic distance between a pair of individuals as a function of allele frequency and kinship coefficient, assuming that SNPs are in Hardy-Weinberg equilibrium.

We also used KING^26^ to identify 30 unrelated individuals from the following 1000 Genomes Project^24^ populations: Yoruba (YRI), CEPH-European (CEU), Finnish (FIN), Colombian (CLM), Peruvian (PEL), Puerto Rican (PUR), and Mexican from Los Angeles (MXL). We used these 30 unrelated individuals per population for all analyses unless otherwise stated (**Supplementary Figure 1**).

### Genotype Data Processing

We generated a joint variant call file (VCF) containing single nucleotide polymorphisms (SNP) from two separate data sets. The first data set contained 210 whole genome sequences sampled from the aforementioned 1000 Genomes Project populations^24^. The second data set contained 449 whole genome sequences from Costa Rican and Colombian individuals. Variants in the second data set were called following the GATK best practices pipeline^27^ with the HaplotypeCaller of GATK. All multi-allelic SNVs and variants that failed Variant Quality Score Recalibration were removed. Genotypes with genotype quality score ≤ 20 were set to missing. Further quality control on variants was performed using a logistic regression model that was trained to predict the probability of each variant having good or poor sequencing quality. Individuals with poor sequencing quality and possible sample mix-ups were removed, and all sequenced individuals had high genotype concordance rate between whole genome sequences and genotypes from microarray data. All sequenced individuals had consistency between the reported sex and sex determined from X chromosome and also between empirical estimates of kinship and theoretical estimates. More information on sequencing and quality control procedures is discussed in Sul et al. 2018 (unpublished data).

We used the following protocol to merge these two datasets. First, we used guidelines from the 1000 Genomes Project strict mask to filter the Costa Rican and Colombian VCFs as well as the 1000 Genomes Project VCFs. Then, we used GATK to remove sites from both sets of VCFs that were not bi-allelic SNPs or monomorphic. Next, we merged the 1000 Genomes Project VCFs with the Costa Rican and Colombian VCFs into a single joint-VCF for each chromosome. We only used autosomes for our analyses. Lastly, we filtered the merged joint-VCF to only contain sites that were present in at least 90% of individuals. There were a total of 57,597,196 SNPs and 1,891,453,144 monomorphic sites in the final data set. We ensured that the merged data sets were comparable by examining the number of derived putatively neutral alleles across the 30 unrelated individuals in all sampled populations, and finding few differences between populations, which is consistent with theory^8^ (**Supplementary Figure 2**).

### Calculating Genetic Diversity

We computed two measures of genetic diversity from sites called across all 30 unrelated individuals from each population: pi (π) and Watterson’s Theta (θ_w_). The average number of pairwise differences per site (π) was calculated across the genome as:

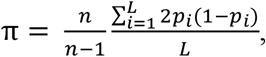

where *n* is the total number of chromosomes sampled, *p* is the frequency of a given allele, and *L* is the length in base pairs of the sampled region. Watterson’s Theta, was computed by counting the number of segregating sites and dividing by Watterson’s constant, or the *n*-1 harmonic number^28^.

### Site Frequency Spectrum (SFS)

Site frequency spectra were generated using the 30 unrelated individuals from each population, SNPs with missing data were removed from these analyses. There was a total of 16 SNPs out of the 57,597,196 SNPs that were removed due to missing data.

### Linkage Disequilibrium Decay

We calculated LD between pairs of SNPs for all unrelated individuals. First, we applied a filter to remove sites that were not at a frequency of at least 10% across all populations. Next, pairwise *r*^2^ values were calculated using VCFTools^29^. SNP pairs were then binned according to physical distance (bp) between each other and *r*^2^ was averaged within each bin.

### Identifying Identity by Descent Segments

To detect regions of the genome that have shared IBD segments between pairs of individuals, we first removed singleton SNPs in each population since singletons are not informative about shared IBD. Then, we called IBD segments using IBDSeq^30^. IBDSeq is a likelihood-based method that is designed to detect IBD segments in unphased sequence data. We chose to use IBDSeq because other methods that require computational phasing could be biased when applied to Latin American population isolates, as they do not have a publicly available reference population to aid in phasing. We compared IBDSeq to two well-known phasing methods Beagle^31^ and GERMLINE^32^ to determine whether it was feasible to use IBDSeq on an admixed population (**Supplementary Figure 3**). Beagle^33^ produced the shortest IBD segments while GERMLINE produced the longest IBD segments. IBDSeq produced segments with a length distribution similar to what we observed in Beagle, though the average segment length was slightly larger, which we expected given that IBDSeq was created to call longer segments that would have previously been broken up when using Beagle for phasing^30^. We used the default parameters for IBDSeq.

Next, we filtered the pooled IBD segments to remove artifacts. First, we calculated the physical distance spanned by each IBD segment. Then, we totaled the number of SNPs that fell within each segment. We observed an appreciable number of IBD segments that were extremely long but sparsely covered by SNPs (**Supplementary Figure 4**). IBD segments were removed if the proportion of the IBD segment covered by SNPs was not within one standard deviation of the mean proportion covered across all IBD segments (**Supplementary Figure 4**). Strong deviations from the mean could indicate that the IBD segment spans a region of the genome with low mappability, and we are only calling the SNPs at the outer ends of the segment. Therefore, the true segment length might be much shorter than what is being calculated by IBDSeq. Lastly, we converted from physical distance to genetic distance using the deCODE genetic map^34^.

### Enrichment analyses of IBD segments

To determine whether certain populations contain more IBD segments than others, we followed the IBD score procedure outlined in Nakatsuka and collegues^17^. A population’s IBD score was calculated by computing the total length of all IBD segments between 3 and 20 cM. The score difference is the difference between the query population’s IBD score and the Finnish IBD score. The score ratio is the ratio of each population’s IBD score relative to the Finnish IBD score. The significance of enrichment relative to the Finnish was evaluated using a permutation test for each population, where IBD segment length was held fixed and labels of the two populations were permuted. We recalculated the score on a total of 10,000 permutations to generate a null-distribution of scores for each isolate.

### Estimating Effective Population Size

We used the output files from IBDSeq to estimate the recent effective population size from the 30 unrelated individuals from each sampled population. We estimated effective population size by using the default settings in IBDNe^35^. We set the minimal IBD segment length equal to 2cM since that is the suggested setting when using sequence data.

### Identifying Runs of Homozygosity

Runs of homozygosity were identified for each individual using VCFTools, which implements the procedure from Auton et al. 2009^36^. Next, we examined the number of callable sites that lie within each ROH. We found that there was a bi-modal distribution of coverage for ROH, where some ROH appeared to contain almost no callable sites, while others had much higher coverage. We only kept ROHs that were at least 2Mb in length, which we called long runs of homozygosity, and were at least 60% covered by callable sites. (**Supplementary Figure 5**).

### Calculating Inbreeding Coefficients

SNP-based inbreeding coefficients were calculated using VCFTools^29^. VCFTools calculates the inbreeding coefficient *F* per individual using the equation 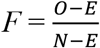, where *O* is the observed number of homozygotes, *E* is the expected number of homozygotes (given population allele frequency), and *N* is the total number of genotyped loci.

Pedigree-based inbreeding coefficients were computed using the R package kinship2^37^.

### Demographic Simulations

In order to investigate how aspects of the population history affect current day genetic diversity in Latin American isolate populations, we simulated genetic variation data using the forward simulation software SLiM 4.2.2^38^. We simulated a sequence length of 10Mb under uniform recombination rate of 1×10^−8^ crossing-over events per chromosome per base position per generation and under a mutation rate of 1.5×10^−8^ mutations per chromosome per base position per generation. Every simulation contained intergenic, intronic, and exonic regions, but only nonsynonymous new mutations experienced natural selection in accordance with the distribution of selection coefficients estimated in Kim et al. 2017^39^. Within coding sequences, we set nonsynonymous and synonymous mutations to occur at a ratio of 2.31:1^39,40^. The chromosomal structure of each simulation was randomly generated, following the specification in the SLiM 4.2.2 manual (7.3), which is modeled after the distribution of intron and exon lengths in Deutsch and Long^41^.

We assumed an effective population size in the ancestral African population of 10,000 individuals, and a reduction in size to 2,000 individuals, starting 50,000 years ago, reflecting the colonization of the European, Asian, and American continents. The population then recovers to a size of 10,000 individuals 5,000 years ago. The colonization bottleneck is assumed to occur 500 years ago by an admixture event with a European population (70% admixture proportion) and is followed by an immediate reduction in population size to 1,000 individuals. The recent expansion in population size is modeled by an increase in population size to 10,000 individuals 200 years ago. We simulated data with recent inbreeding and without recent inbreeding. In the former case, inbreeding started at the time of the European colonization 500 years ago and continues until the present. Inbreeding is implemented with the “mateChoice” function in SLiM. Because SLiM’s pedigree track function is only valid for at most second-degree related individuals, 50% of the time, mating occurs randomly. However, in the remaining cases, mating occurs between close relatives with a relatedness coefficient bigger than 0.25. This produces levels of consanguinity similar to those seen empirically as measured by F (see Results). Finally, we sampled a total of 60 random individuals and calculated summary statistics on the sample data. The simulation script can be found on GitHub (see Web Resources).

### Annotation of Variants

The ancestral allele was determined using the 6-primate EPO alignment (see Web Resources) and we restricted to only those sites called with the highest confidence. After filtering, 54,049,081 SNPs remained.

Subsequently, exonic SNPs were annotated using the SeattleSeq Annotation website (see Web Resources). A total of 693,301 SNPs were annotated as either nonsynonymous or synonymous. We further classified these sites as either putatively neutral or deleterious using Genomic Evolutionary Rate Profiling (GERP) scores^42^. A GERP score less than two was considered as putatively neutral and a GERP score greater than 4 was considered as putatively deleterious, totaling 404,302 classified SNPs.

### Counting Deleterious Variants

We used three different statistics to count the number of deleterious mutations per individual. First, we tabulated the number of deleterious variants (the number of heterozygous plus the number homozygous derived genotypes). Second, we counted the total number of derived deleterious alleles (the number of heterozygous genotypes plus twice the number of homozygous derived genotypes). Third, we computed the total number of derived deleterious homozygous genotypes.

### Testing for an enrichment of deleterious variation in ROHs

We were interested in whether there is an enrichment of nonsynonymous or loss-of-function mutations in ROH over non-ROH regions for the three different ways of counting deleterious variants outlined above. To account for differences in neutral variation, we standardized by synonymous variation, which is assumed to be neutral. Then, we calculated the ratio of nonsynonymous over synonymous variation in ROH regions divided by the ratio of nonsynonymous over synonymous variation outside of ROH. We computed significance using a permutation test, where the position of each SNP and its annotation as synonymous versus nonsynonymous was fixed and the positions of the vector of ROH annotations were randomly placed throughout the genome. Thus, the frequency distribution of synonymous and nonsynonymous SNPs, as well as the total amount of ROH and non-ROH annotations, is kept constant when compared to the unpermuted data. We recalculated the ratio for a total of 10,000 permutations to form a null-distribution of ratios and then computed significance.

### Calculating Ancestry Proportions

We estimated genome-wide ancestry proportions in members of the CR and CO pedigrees using ADMIXTURE^43^ (v1.22). Then the genome-wide ancestry proportions from ADMIXTURE were used as the prior in local ancestry analysis using LAMP^44^. We generated estimates for all 838 pedigree members with SNP array genotype data. Detailed information on the SNP array data can be found in Pagani et al.^45^. The reference populations were the CEU (n=112) and YRI (n=113) from HapMap^46,47^, as well as 52 Native American samples from Central or South America. The Native American samples are the Chibchan-speaking subset of those used in Reich et al.^48^, selected to originate from geographical regions relevant to CR/CO and to have virtually no European or African admixture. In total the admixture analysis used 57,180 LD-pruned SNPs and 1115 individuals.

### Accounting for Relatedness

We tested for correlations among several quantities computed for each individual in the Latin American population isolates. Because some of these individuals are closely related, the data points in our linear regression are no longer independent. Therefore, we implemented the R-package GenABEL^49^ to incorporate kinship when performing statistical tests for our correlations. We used the polygenic_hglm() function where the *formula* input was the equation for our linear model of interest and the *kinship.matrix* input was a kinship matrix computed from our pedigree computed using kinship2^37^. Our input took the following form: (F_PED_ ~ Length of genome in ROH, kin = kinshipMatrix, data = df).

## Results

### Genetic Variation in Population Isolates

We first compared levels of genetic diversity in a sample of 30 unrelated individuals across the 1000 Genomes populations^24^ and the CO and CR isolates. We split the genome into several different genomic regions and in each region summarized genetic variation using both the average number of pairwise differences (π) and Watterson’s theta (θ_w_) (**Figure 1A and B**). Overall, we found differences in diversity across the functional category of sequence studied in all populations, with coding regions exhibiting the lowest diversity and intergenic regions the highest. These patterns are consistent with the role of purifying selection affecting coding diversity^39^. However, if we look genome-wide or focus on intronic regions we see intermediate levels of diversity (**Supplementary Table 1 and 2**). We suspect that these categories are more strongly influenced by linked selection^50–52^.

**Figure 1.**
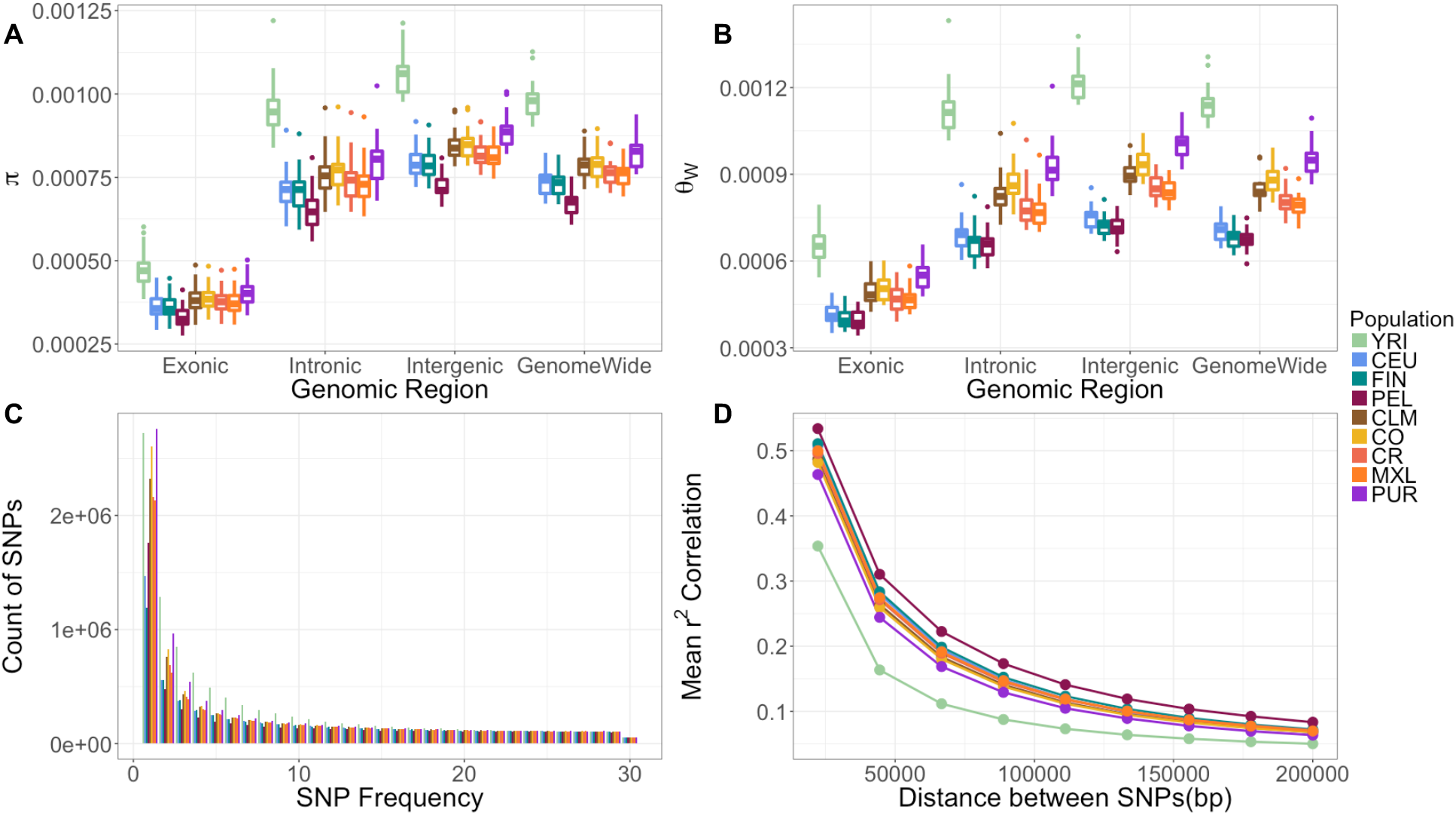
Patterns of genetic variation in the Colombian and Costa Rican populations compared to the 1000 Genomes populations. (A) Diversity measured using the average pairwise differences between sequences, π. (B) Diversity measured using the number segregating sites, Watterson’s theta (θ_W_). (C) The site frequency spectrum for each population. (D) Average LD (*r*^2^) between pairs of SNPs. All statistics were calculated using 30 unrelated individuals per population (see Methods). Box plots in (A) and (B) show the distribution over 22 autosomes. YRI: Yoruba 1000 Genomes; CEU: Ceph-European 1000 Genomes; FIN: Finnish 1000 Genomes; PEL: Peruvian 1000 Genomes; CLM: Colombian 1000 Genomes; CO: Colombia; CR: Costa Rica; MXL: Mexican from Los Angeles 1000 Genomes; and PUR: Puerto Rican 1000 Genomes.

As we are interested in the role of demography in shaping genetic diversity, we focused on comparisons of intergenic levels of diversity as those are most likely to be neutrally evolving (**Figure 1A and B**). Overall, the YRI had the highest level of diversity (π ≈ 0.0010; θ_w_ ≈ 0.0012) (**Supplementary Table 1 and 2**). The European populations (CEU and FIN) had lower levels of diversity. The CEU and FIN had similar levels of π (approximately 0.0004), despite the FIN being considered an isolated population. However, the FIN had reduced numbers of SNPs as reflected by lower values of θ_w_ (CEU ≈ 0.0008 & FIN ≈ 0.0008). The CO and CR had levels of diversity similar to that of several other Latin American populations in the 1000 Genomes Project (CLM and MXL). We found no clear pattern of the population isolates (FIN, CO, CR) having lower diversity than their most similar non-isolated population. Instead, diversity levels tended to be higher across all the sampled Latin American populations (CLM, CO, CR, MXL, and PUR) when compared to the European populations. One exception to this pattern is the PEL population, who had the lowest neutral levels of diversity (π ≈ 0.0007; θ_w_ ≈ 0.0007).

Next, we examined the proportional site frequency spectrum (SFS; **Figure 1C**). Latin American populations had the highest proportion of singletons. The CO and CR had similar proportions of singletons when compared to other 1000 Genomes Project Latin American populations. Conversely, the FIN had the lowest proportion of singletons in comparison to all sampled populations. The depletion of singletons relative to common variation supports the presence of a stronger founder effect during the FIN population history^13^.

We also examined patterns of linkage disequilibrium (LD), since LD is affected by population size and recent bottlenecks^53,54^. **Figure 1D** shows the mean decay of *r^2^* with physical distance over 2Mb intervals across the genome in each population. We found that the YRI had the lowest levels of LD for each bin of physical distance, and the PEL formed the upper bound of the LD decay curves. The remaining Latin American populations (PUR, MXL, CLM, CO, CR) clustered together, close to the YRI, while the CEU and FIN are shifted toward higher values, like those seen in the PEL.

The FIN were previously shown to have more extensive haplotype blocks in their genome in comparison to the Latin American isolates^9^. In line with these findings, we observed faster LD decay in the Latin American isolates relative to the FIN. When considering pairs of SNPs 150kb or more apart, rates of LD decay become quite similar across all the sampled populations. Analogous to other diversity statistics, LD in the CO and CR closely resembled those of non-isolated Latin American populations. Once again, we found there is no clear pattern of having lower diversity or more LD that holds across all the population isolates (FIN, CO, CR) when compared to their most similar non-isolated population.

### Latin American isolates carry more IBD segments than Finnish

Next, we used IBD sharing between pairs of individuals to gain insight about more recent demographic events within populations (**Figure 2**). We compared the amount of IBD within each population by computing an IBD score. Each population’s IBD score was calculated by totaling the length of IBD segments between 3cM and 20cM. We expressed IBD scores for each population as the ratio of the IBD score for a given population relative to the IBD score in the FIN (**Figure 2A**). We also tabulated the total count of IBD segments for each population. The CEU showed the lowest number of both called IBD segments and the lowest IBD score relative to the FIN (p-value = 0.0001). Latin American populations formed the upper bounds of both total IBD segments called and IBD enrichment scores (**Figure 2A**). The PUR had the largest number of IBD segments (1402) and had a 2.1-fold increase in IBD score relative to the FIN (p-value < 1×10^−4^). The CO and CR isolates had a 1.8-fold and 2-fold increase in their IBD scores relative to the FIN (p-value < 1×10^−4^), as well as carrying more IBD segments than the FIN (**Figure 2B and 2C**). However, there were some Latin American populations that exhibited depletions in both IBD segments and IBD scores relative to the FIN. The MXL and PEL have the lowest number of IBD segments for the Latin American populations. Previous work has shown that a larger effective population size in admixed populations likely drove the depletion of IBD segments in these two Latin American populations^55^.

**Figure 2.**
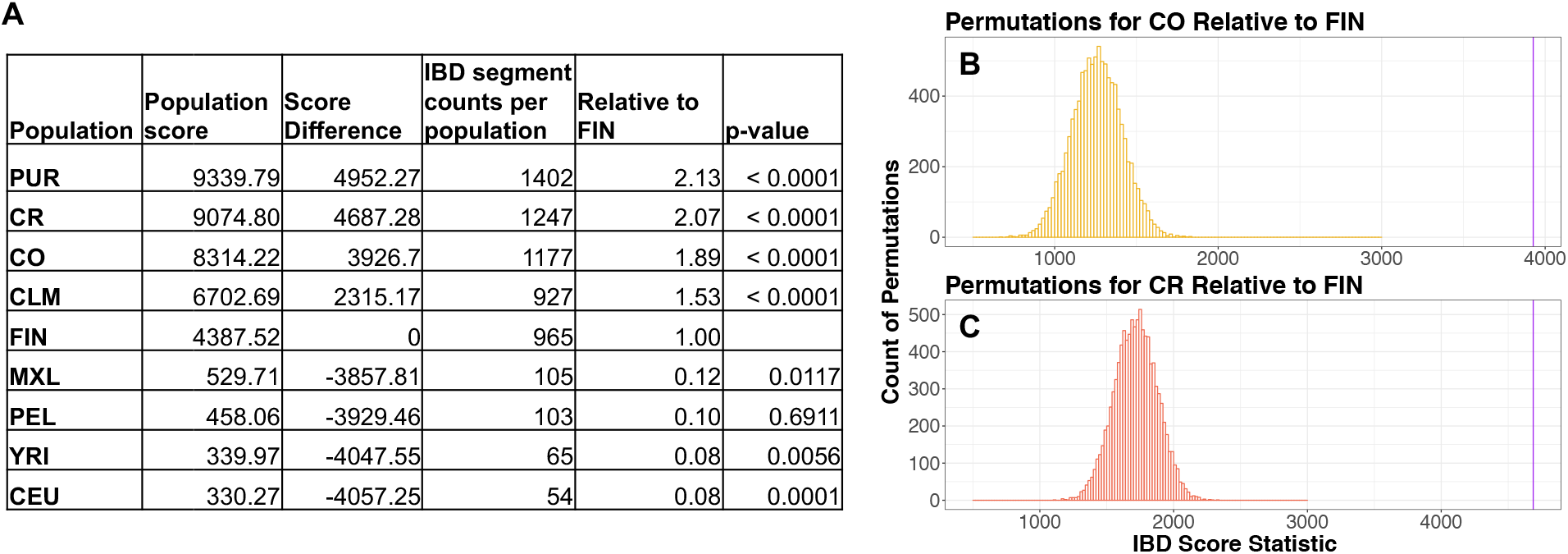
Latin American population isolates (CR and CO) have significantly more identity by descent (IBD) segments relative to the Finnish (FIN). IBDSeq was used to generate IBD segments for the 30 unrelated individuals in each population. (A) Population score was calculated by summing all IBD segments between 3cM and 20cM for each population. Score difference is the population score minus the FIN population score. IBD enrichment for each population score is reported as relative to the FIN (i.e. FIN score is 1.0). (B&C) Histogram of 10,000 permutation tests of Colombia (p < 1.0 e^-04^) and Costa Rica (p < 1.0 e^-04^) population scores versus Finnish score. The observed score for each population is demarcated by the purple line. Population abbreviations are as in **Figure 1.**

### Inferring the Demographic History of Latin American Isolates

We next leveraged the patterns of IBD described above to estimate the effective population size using IBDNe^35^ on the 30 unrelated individuals from each population (**Figure 3**). The use of only 30 unrelated individuals caused limitations for accurate estimation of N_e_ (see Discussion), but the demographic history of the population was robust to the number of individuals used. First, we found that recent demography differs vastly between the European populations (FIN and CEU). In general, CEU experienced population expansions over much of their demographic history. It was only in the most recent generations that they experienced a decrease in N_e_. The FIN, on the other hand, have experienced a long population decline since their founding, approximately 4000 years ago, followed by a recent population expansion.

**Figure 3.**
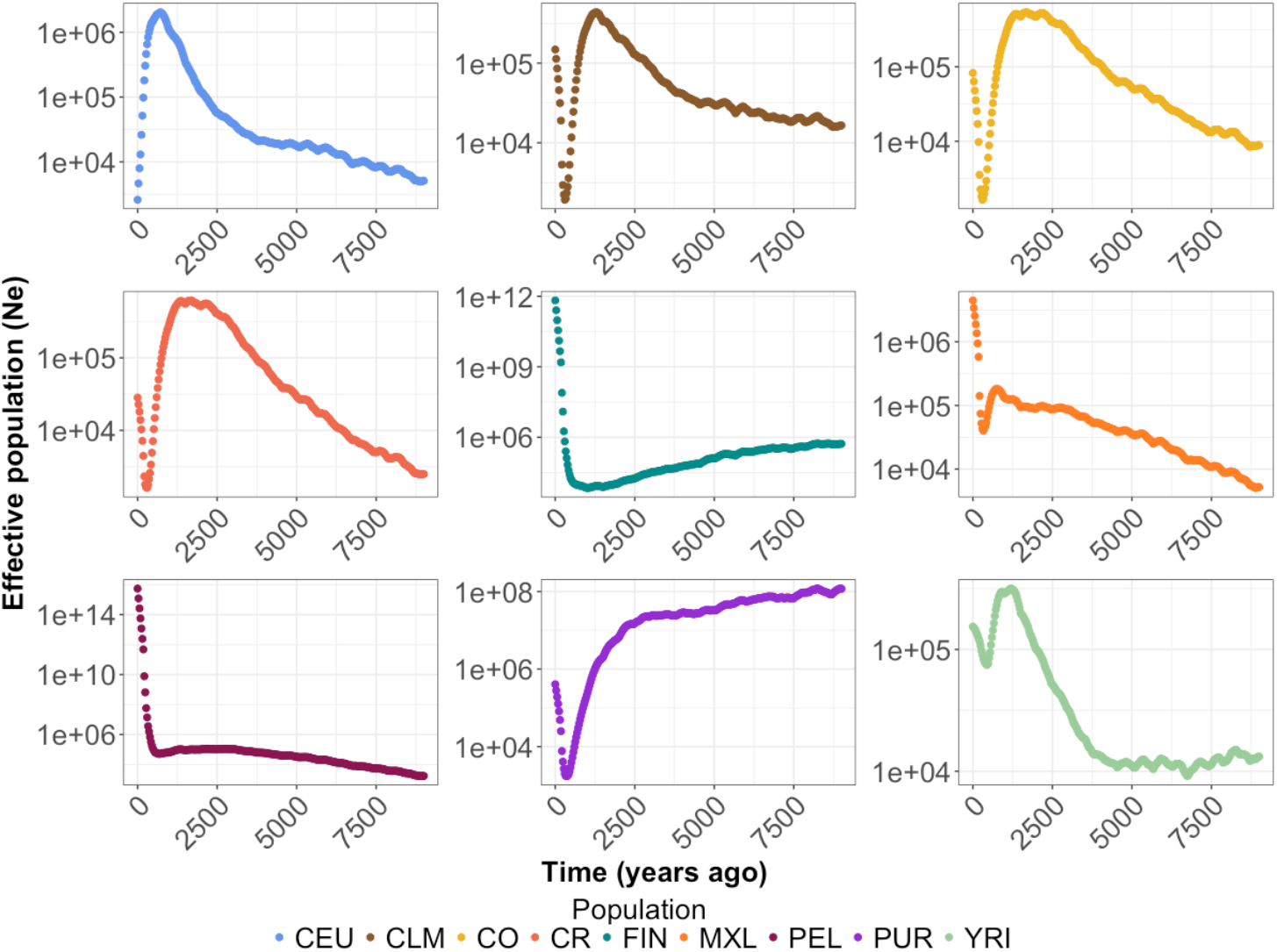
Recent effective population size differs across populations. IBDNe^35^ (see Methods) was used to infer effective population size (Ne) over the last 9000 years for each population. Note the FIN shows a long slow decline followed by recent growth. The CO and CR show sharp bottlenecks approximately 500 years ago followed by recent growth. Population abbreviations are as in **Figure 1.**

When analyzing the Latin American isolates, we detected a recent bottleneck, approximately 500 years ago (**Figure 3**). This bottleneck could correspond to the recorded bottleneck that followed the founding of these populations, and it appears to be much shorter and less severe than the bottleneck seen in the FIN. The strength and duration of bottlenecks varied across each of the Latin American populations. For example, we observed a more severe bottleneck in the CR, CO, CLM, and PUR than in PEL or MXL. However, we detected a subsequent period of growth across all populations following the bottleneck. The rate of growth differed across each population, and the PEL appeared to be growing at a much more rapid rate than any of the other Latin American populations.

### Exploring Recent Consanguinity

Isolated populations may have experienced recent consanguinity. To test for this, we began by examining SNP-based inbreeding coefficients (F_SNP_) (**Supplementary Figure 6**). YRI individuals had the lowest median inbreeding coefficients and the CO and CR isolates had the highest median inbreeding coefficients. Further, the CO and CR also had the highest maximum F_SNP_ values in the entire sample of unrelated individuals from any population (**Supplementary Figure 6**). Median levels of F_SNP_ in the CEU suggested that they are more inbred than the FIN, which may be a result of how 1000 Genomes samples were selected. The PEL had the largest variance in F_SNP_ across any of the sampled populations.

Next, we examined patterns of long runs (>2Mb, see Methods) of homozygosity (ROH), since ROH have been linked to recent consanguinity^20,21,56–58^. The YRI and CEU had the lowest amount of their genome contained within an ROH (**Figure 4A**). The FIN had higher median amounts of their genome within an ROH in comparison to the CEU. Latin American isolates had the highest median amount of the genome contained within an ROH. Specifically, the CR had the highest median at 21.7 Mb. Further, the Latin American isolates also had the greatest variance in the amount of the genome contained within an ROH. For example, one of the CO individuals had approximately 230 Mb of her/his genome contained in long ROHs.

**Figure 4.**
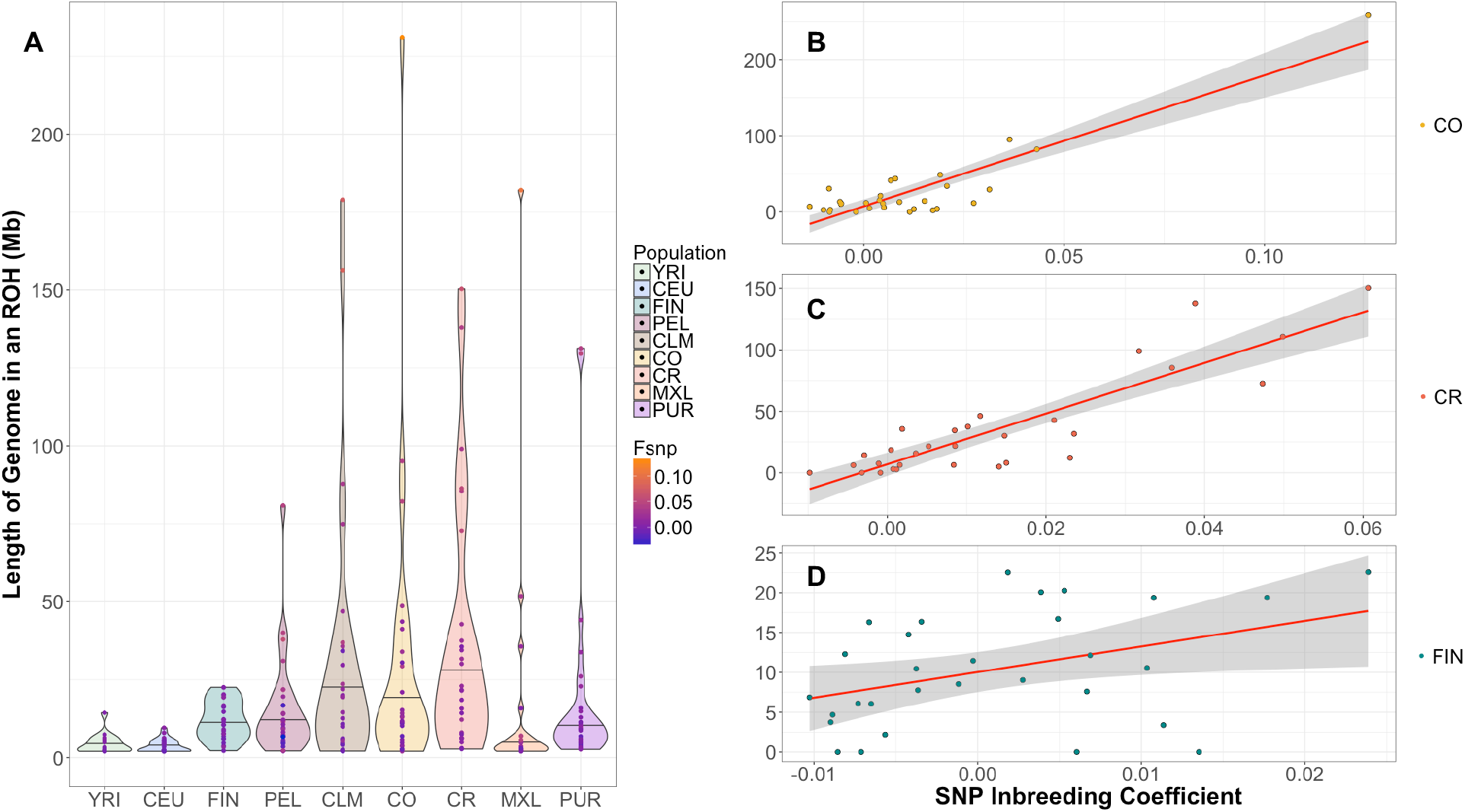
Length of the genome in a run of homozygosity (ROH) varies across populations and correlates with SNP inbreeding coefficient. The length of the genome in an ROH was calculated for each unrelated individual (*n*=30 per population) by summing the physical distance (Mb) of each ROH >2Mb. (A) The length of the genome in an ROH varies by population. The black line within the violin marks the median. F_SNP_ for each individual was overlaid within the ROH violin plot. A blue hue indicates the lowest F_SNP_ and orange indicates the highest F_SNP_. (B) Length of the genome in an ROH is strongly correlated with F_SNP_ in Colombians, (*R*^2^ = 0.8060, p –value = 1.1 × 10^−11^). (C) Length of the genome in an ROH is strongly correlated with F_SNP_ in Costa Ricans, (*R*^2^ = 0.7740, p-value = 9.5 × 10^−11^). (D) Length of the genome in an ROH is positively correlated with F_SNP_ in Finnish, (*R*^2^ = 0.1288, p-value = 0.03). Population abbreviations are as in **Figure 1**.

As expected, we found that the amount of the genome contained in a long ROH strongly correlated with an individual’s F_SNP_ (CO: *R*^2^ = 0.8060, p –value = 1.1 × 10^−11^; CR: *R*^2^ = 0.7740, p-value = 9.5 × 10^−11^; FIN: *R*^2^ = 0.1288, p-value = 0.03) (**Figure 4B-4D**). Indeed, individuals with higher values of F_SNP_ tended to have more of their genome within an ROH. Further, the individual with the highest F_SNP_ (0.133) also had the largest amount of his/her genome in long ROHs (230Mb).

The total number of ROH segments per individual followed a similar pattern as the total amount of genome within an ROH (**Supplementary Figure 7**). For example, in populations with low values of F_SNP_, ROH segments were not frequent. One YRI individual and three CEU individuals carried a ROH >4Mb, whereas more than 50% of CO and CR individuals carried an ROH >4Mb. Additionally, the longest ROHs identified (>20MB), only occurred in Latin American populations, where there were the largest values of F_SNP_ (**Supplementary Figure 7**).

Importantly, the FIN individuals had significantly fewer ROH segments than the CO and CR, and most individuals had an F_SNP_ close to 0 (**Figure 4**). The Latin American isolates had the most ROH in comparison to any other sampled population, as well as the largest values of F_SNP_ (**Figure 4**).

### Determining the Mechanisms that Generate Runs of Homozygosity

In principle, ROHs can be generated either by recent consanguinity over the last few generations, or by older historical processes, such as bottlenecks^19,56,58–60^. Based on both historical data^23^ and inference from IBDNe analyses, Latin American population isolates show evidence of recent population bottlenecks. Therefore, we used two complementary strategies to test whether recent consanguinity or bottlenecks drove the observed increase in ROHs in the Latin American isolates. First, we used the extensive pedigree data for 449 sequenced individuals to calculate a pedigree inbreeding coefficient (F_PED_) for each individual (**Figure 5**). Most individuals had a F_PED_ of 0. However, there were several individuals with values of F_PED_ as high as 0.07 in CR and 0.06 in CO. We observed a significant correlation between F_SNP_ and F_PED_ (R^2^=0.1520 and p-value < 2 x10^−16^), even after accounting for the non-independence of individuals based on their kinship (**Figure 5A**; see Methods). These correlations suggest that the recent consanguinity captured within the last few generations in the pedigree was likely sufficient to drive the increase in ROHs in the CO and CR populations. F_SNP_ was a substantially better predictor of the amount of an individual’s genome that falls within a ROH (R^2^ = 0.7540 and p-value < 2 × 10^−16^), than F_PED_ (R^2^ = 0.2180 and p-value < 2 × 10^−16^) (**Figure 5B and C**) likely due to the fact that F_SNP_ captured distant background relatedness within the population as well as the realized level of consanguinity, rather than the expected value^61^. Further, because the pedigrees were ascertained and analyzed separately, connections between pedigrees were not accounted for in F_PED_, but were likely captured by F_SNP_.

**Figure 5.**
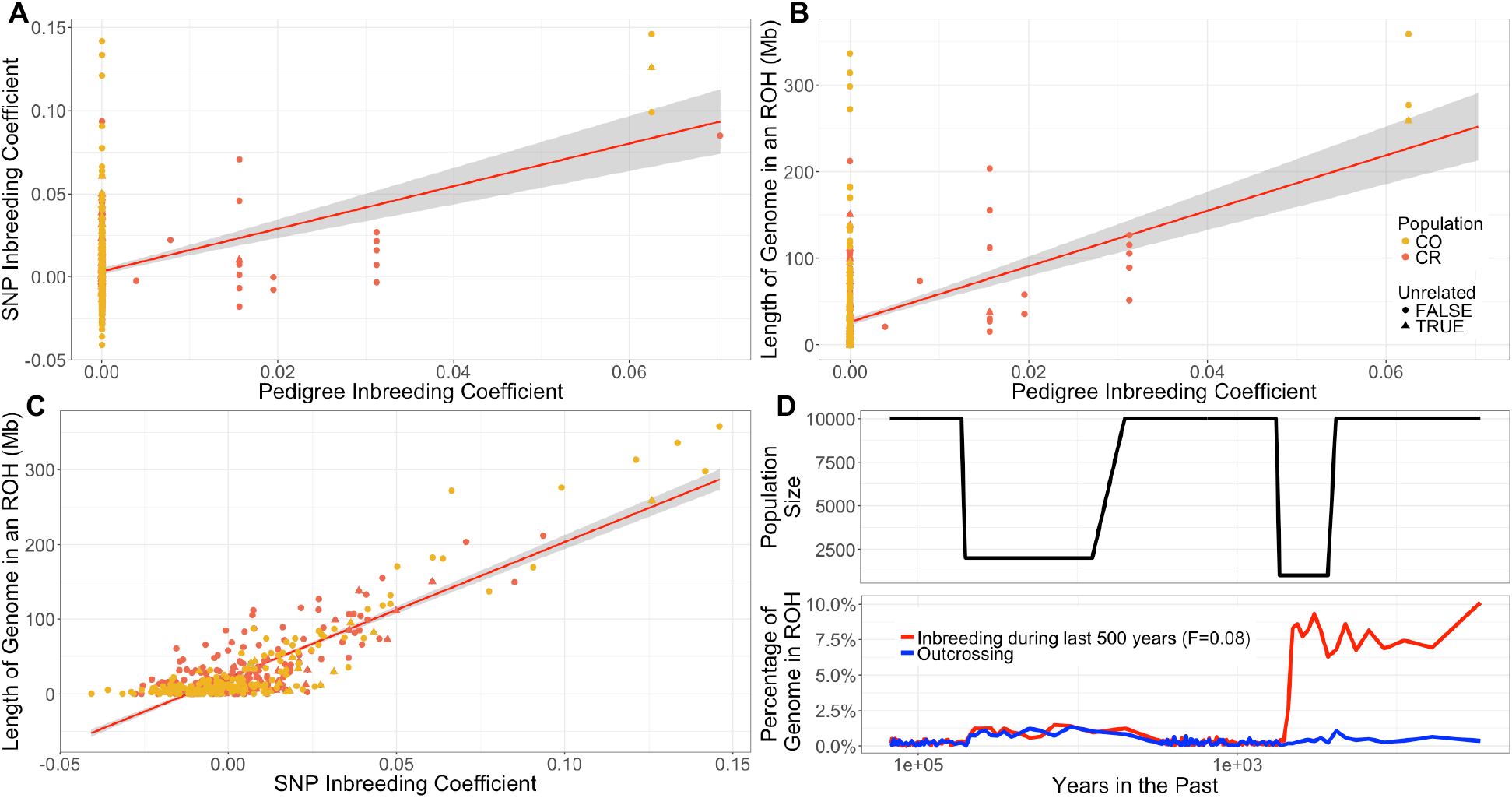
Recent consanguinity creates ROH in Costa Rica and Colombia. Triangles represent the individuals that were sampled in the unrelated data set (*n*=30). (A) F_SNP_ is correlated with the pedigree inbreeding coefficient (F_PED_; *R*^2^=0.1520, p-value < 2 x10^−16^) in the full data. (B) The length of the genome in an ROH is correlated with F_PED_ (*R*^2^ = 0.2180, p-value < 2 × 10^−16^). (C) The length of the genome in an ROH is correlated with F_SNP_ (*R*^2^ = 0.7540, p-value < 2 × 10^−16^). (D) Forward simulations show that recent consanguinity during the last 500 years can generate ROHs while bottlenecks cannot. Top panel shows the changes in population size used in the simulations. Bottom panel shows how the percent of the simulated in genome within an ROH changes over time. Population abbreviations are as in **Figure 1**.

As a second approach to determine the mechanism driving the increase in ROHs in the CO and CR populations, we conducted forward in time demographic simulations. We simulated a 10Mb region under a demographic model that reflected changes in effective population size during the human expansion across the European, Asian and American continents, as well as the more recent bottleneck during the Spanish colonization about 500 years ago (**Figure 5D**; see Methods). We compared simulations with no inbreeding to simulations with recent inbreeding. Consanguinity in the populations was modeled to begin 500 years ago, and simulated individuals had an inbreeding coefficient of about 0.075. This level of inbreeding was comparable to the level of inbreeding in some of the CO and CR individuals, based on calculations using pedigree data (see Methods). Our simulations suggested that the recent population bottleneck caused by the Spanish colonization was not capable of generating the large amounts of the genome within an ROH (>2Mb) that we observed for some of the individuals (**Figure 5D**). Only when simulating recent inbreeding could levels of the genome in an ROH comparable to that we observed be generated. Thus, recent inbreeding was paramount for generating the long ROH that we observed in the CO and CR isolates.

### Local Ancestry

Since the Latin American isolates originated from an admixture event between Native Americans, Africans, and Europeans, we tested for a correlation between F_PED_ and the proportion of European, African, or Native American ancestry (**Supplementary Figure 8**). We used the entire sequenced Costa Rican and Colombian data set (n=449) for the local ancestry analyses and accounted for relatedness of individuals in all the following reported p-values (see Methods). We found that European ancestry was positively correlated with F_PED_ (p-value = 0.0052) while Native American ancestry was negatively correlated with F_PED_ (p-value = 0.0245). African ancestry was also negatively correlated with F_PED_ (p-value = 0.0496).

Then, we asked if the proportion of ancestry was correlated with the amount of the genome within an ROH (**Supplementary Figure 8**). The correlation between ancestry and amount of the genome within an ROH followed the same trend as the correlation between ancestry and F_PED_. Native American ancestry and African ancestry are negatively correlated with the amount of the genome within a long ROH (p-value = 3.91 × 10^−14^ and p-value = 6.76 × 10^−07^, respectively). European ancestry was positively correlated with the amount of an individual’s genome within an ROH (p-value < 2 × 10^−16^).

### Recent Consanguinity is Correlated with an Increase of Deleterious Variation

It is well known that demography impacts patterns of deleterious variation in populations^5,8,20,62–66^. Thus, we compared patterns of putatively deleterious variation in the CO and CR to those in the FIN. Variants were classified as putatively deleterious or putatively neutral using GERP scores (see Methods). Recall that we consider three ways of counting deleterious variants in the genome of an individual: first, counting the number of heterozygous genotypes plus twice the number of homozygous derived genotypes (i.e. the total number of derived deleterious alleles), second, counting the number of heterozygous and homozygous derived genotypes (counting variants), and third, counting only the number of homozygote derived genotypes (counting homozygotes). The first quantity is most relevant if deleterious alleles are additive, while the third is most relevant if they are recessive. First, we looked at absolute counts of derived deleterious variation across isolates (**Supplementary Figure 9**). Then, we used linear regression to test if there was a relationship between the amount of an individuals’ genome in an ROH and the number of nonsynonymous sites in the genome for each counting method (**Figure 6**).

**Figure 6.**
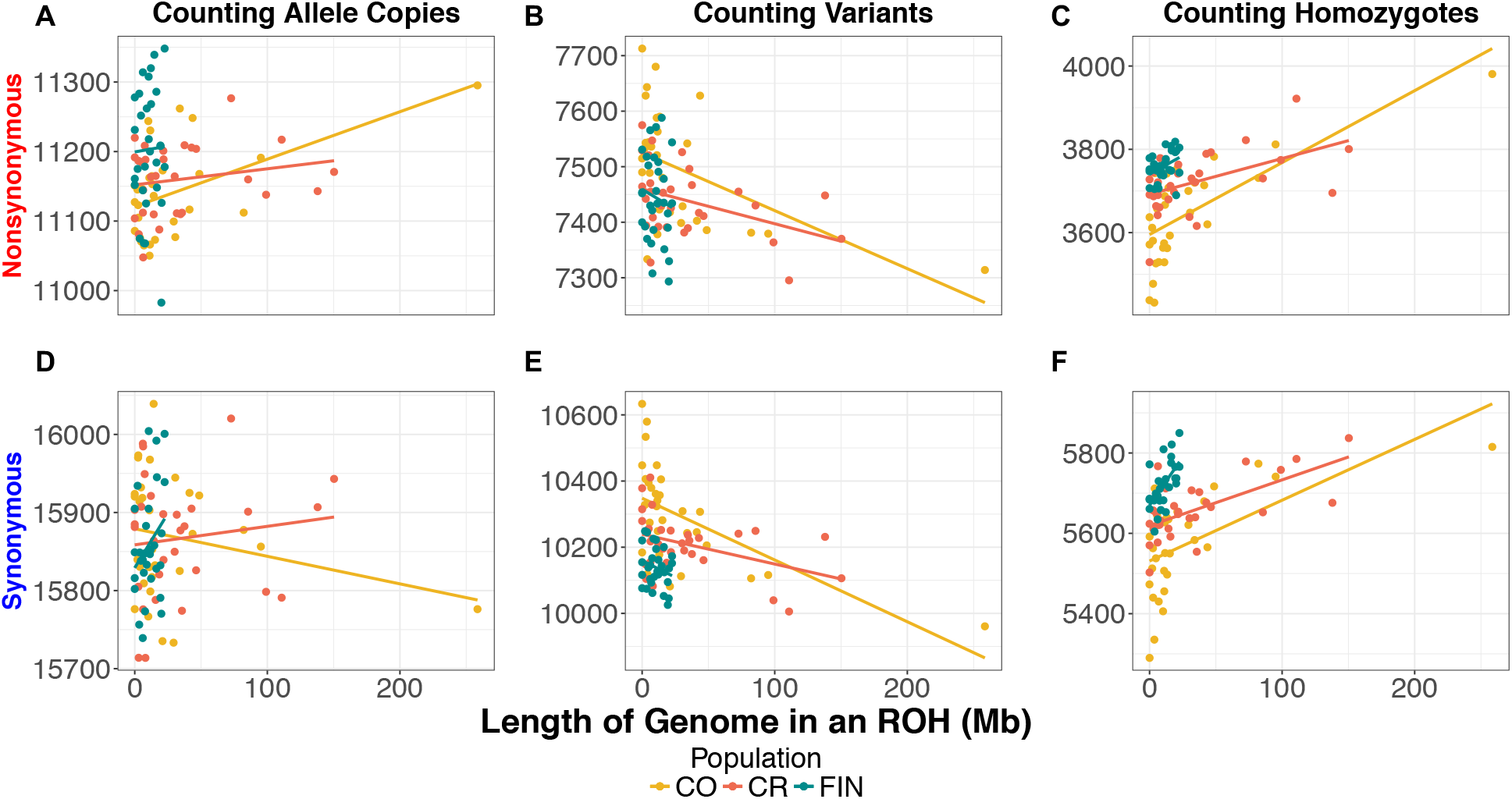
The correlation between ROH and nonsynonymous variation in the Colombian, Costa Rican, and Finnish samples. The count of nonsynonymous and synonymous mutations per individual as a function of the length of the genome in an ROH in the Colombia (CO), Costa Rican (CR) and Finnish (FIN) populations: (A) Number of nonsynonymous alleles per individual. (B) Number of nonsynonymous variants per individual. (C) Number of homozygous nonsynonymous genotypes per individual. (D) The number of synonymous alleles per individual. (E) The number of synonymous variants per individual. (F) The number of homozygous synonymous genotypes per individual. Population abbreviations are as in **Figure 1**.

The FIN carried approximately 1% more derived deleterious nonsynonymous alleles per individual than CO and CR (p-value = 0.0007; p-value = 0.0013). However, there was no significant difference in the number of putatively neutral synonymous derived alleles per individual. These results suggest that the difference seen for putatively deleterious variants is not driven by data artifacts (**Supplementary Figure 9**), and the FIN indeed have a slightly higher additive genetic load than the CO or CR. Turning to the number of variants per individual, FIN individuals carried significantly more deleterious nonsynonymous variants than the CR but not the CO (p-value = 0.0110). However, CO and CR did not differ significantly in the number of deleterious variants carried per individual (**Supplementary Figure 9**). When we examined neutral synonymous variants, CO had significantly more variants than either FIN or CR (p-value = 8.56^-06^; p-value = 0.0054, respectively). Finally, when counting the number of homozygous derived genotypes, we found that the FIN carried 3.3% more deleterious variants in the homozygous state per individual than CO but not the CR (p-value = 0.0003) (**Supplementary Figure 9**). Additionally, the FIN carried significantly more neutral homozygous genotypes per individual than either population (CO p-value = 1.01 × 10^−05^; CR p-value = 6.96 × 10^−05^). The increased deleterious and neutral variation in homozygous form is an expected consequence of the long-term bottleneck that the FIN experienced during their founding.

We next tested whether the amount of the genome in an individual contained within a ROH was correlated with the number of nonsynonymous mutations carried by the individual. Counting nonsynonymous (NS) or synonymous (SYN) allele copies did not show any correlation with the amount of an individuals’ genome that falls within an ROH for the CR or FIN (**Figure 6A and D; Supplementary Figures 10-13**). However, in the CO, as the amount of the genome within an ROH increased, individuals tended to carry more NS alleles, though this correlation was strongly driven by a single individual, who also had the highest F_SNP_ and F_PED_ (R^2^ = 0.2393; p-value = 0.0036; **Supplementary Figure 10**). Importantly, the number of SYN alleles per individual was not correlated with the amount of the genome in an ROH (p-value = 0.2261).

When counting variants per individual, we observed a significant negative correlation with the amount of an individuals’ genome that falls within an ROH in the Latin American isolates (**Figure 6B and E; Supplementary Figures 10-12**). The negative correlation is a result of heterozygous sites being lost when an ROH is formed due to inbreeding. Conversely, when counting homozygous genotypes per individual, we observed a significant positive correlation with the amount of an individual’s genome that falls within an ROH in both the Latin American isolates and FIN (**Figure 6C and F; Supplementary Figures 10-13**). Homozygous genotypes were the only statistic that correlated significantly with the amount of the genome in an ROH across all isolated population for both SYN and NS sites. We observed a stronger correlation between the number of NS homozygous genotypes and the amount of an individual’s genome within an ROH in the Latin American isolates (R^2^ = 0.5000 (CO) & R^2^ = 0.2165 (CR); p-value = 7.546^-06^(CO) and p-value = 0.0059(CR)) compared to the FIN (R^2^ = 0.1130 and p-value = 0.0389) (**Supplementary Figures 10-13**). This pattern exists because the majority of CO and CR individuals carried larger proportion of their genome within an ROH while the FIN individuals do not harbor many ROH.

We next asked whether there was an enrichment or depletion of NS variants relative to SYN variants within versus outside of an ROH using a permutation test on the three different counting approaches (see Methods). When variants or allele copies were counted, none of the populations produced significant results (**Table 1**). When homozygous genotypes were counted, ROHs in the MXL and CR were enriched for homozygous NS genotypes relative to SYN homozygous genotypes (p-value = 0.0052 and p-value = 0.0169) (**Table 1**). Additionally, if we pooled the CR and CO populations, we also observed a significant enrichment of deleterious variation within an ROH compared to non-ROH regions of the genome (p-value = 0.0011).

**Table 1.**
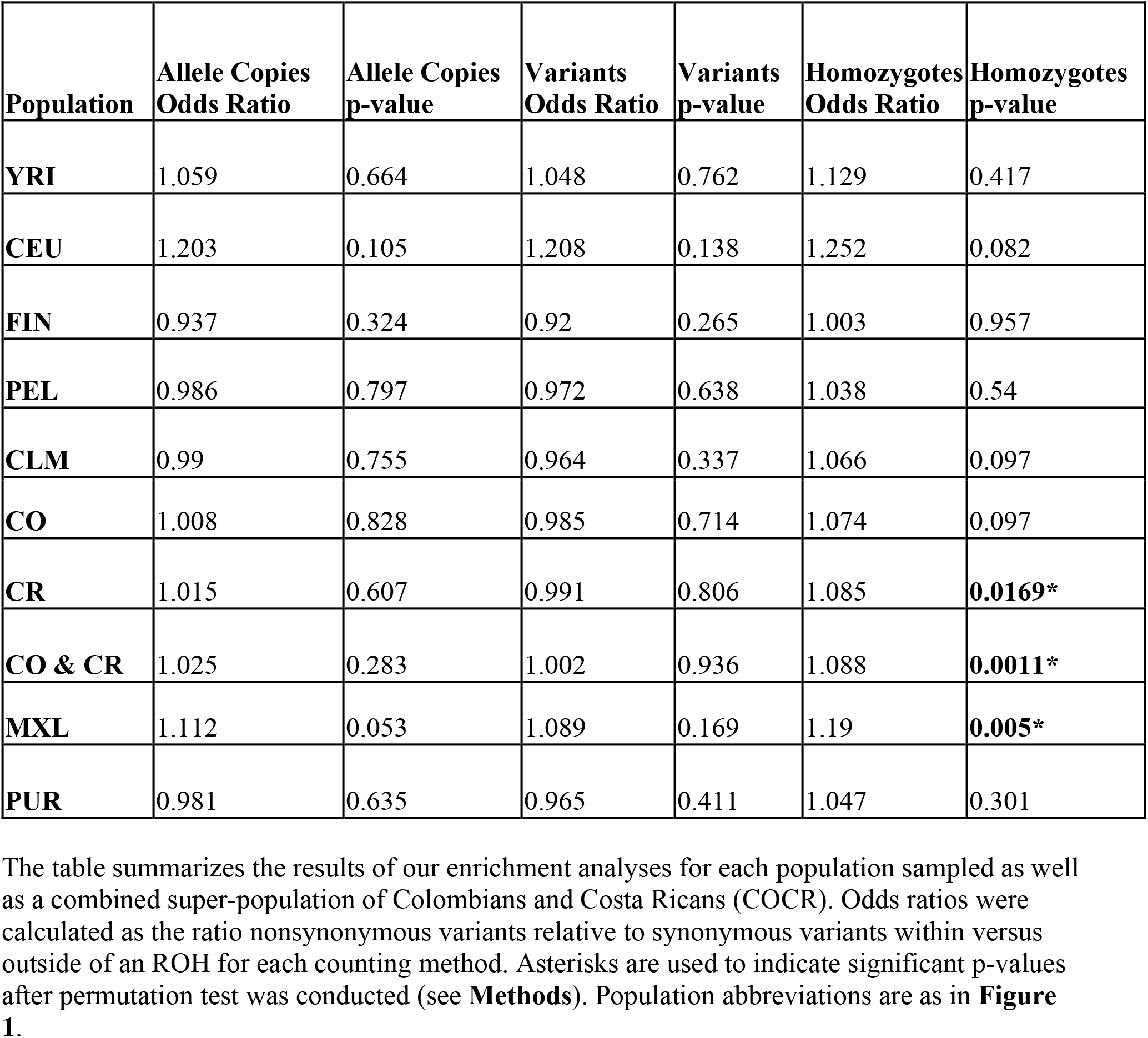
Enrichment of nonsynonymous homozygous derived genotypes within ROHs.

We tested whether F_SNP_ was correlated with the amount of deleterious variation per individual. We only used isolates for these regressions, because we are particularly interested in how recent consanguinity affected deleterious variation in the genome. We observed the exact same pattern with F_SNP_ as with ROH (**Supplementary Figure 14**). Briefly, counting NS or SYN allele copies did not show any correlation with F_SNP_ for the CR or FIN, but there was a significant correlation with NS allele copies in CO (**Supplementary Figure 14; Supplementary Figures 15-17**). Counting NS and SYN variants per individual produced a significant negative correlation with F_SNP_ in the Latin American isolates (**Supplementary Figure 12; Supplementary Figures 15-17**). Counting the number of NS and SYN homozygous genotypes per individual was positively correlated with F_SNP_ in the both Latin American isolates and FIN (**Supplementary Figure 14; Supplementary Figures 15-17**). Again, counting homozygotes was the only method with significant results across all isolated populations for both SYN and NS variants. The ability to recapitulate the pattern we observed in ROH using F_SNP_ was reassuring and adds further support to the strong relationship between recent consanguinity and ROH.

Lastly, because we had multi-generational pedigrees for the Latin American isolates, we examined the correlation between putatively deleterious variation and recent consanguinity as measured by (F_PED_). All the following reported p-values account for kinship (see Methods). When we pooled the CO and CR individuals together, we did not observe any relationship between counting derived deleterious allele copies and F_PED_ after correcting for kinship (**Figure 7A**). Moreover, we observed a negative correlation between F_PED_ and the number of deleterious variants per individual (R^2^ = 0.0375, p-value = 6.02 × 10^−06^). The number of neutral variants per individual was also negatively correlated with F_PED_ (p-value = 2.26 × 10^−10^) (**Figure 7B**). Finally, we observed a positive correlation between F_PED_ and derived deleterious homozygotes (R^2^ = 0.0575, p-value = 1.0 × 10^−06^) as well as between F_PED_ and the number of neutral derived homozygotes per individual. (p-value = 1.03 × 10^−08^) (**Figure 7C**). These results suggest that recent consanguinity during the last few generations has increased the number of derived deleterious homozygous genotypes in these two populations.

**Figure 7.**
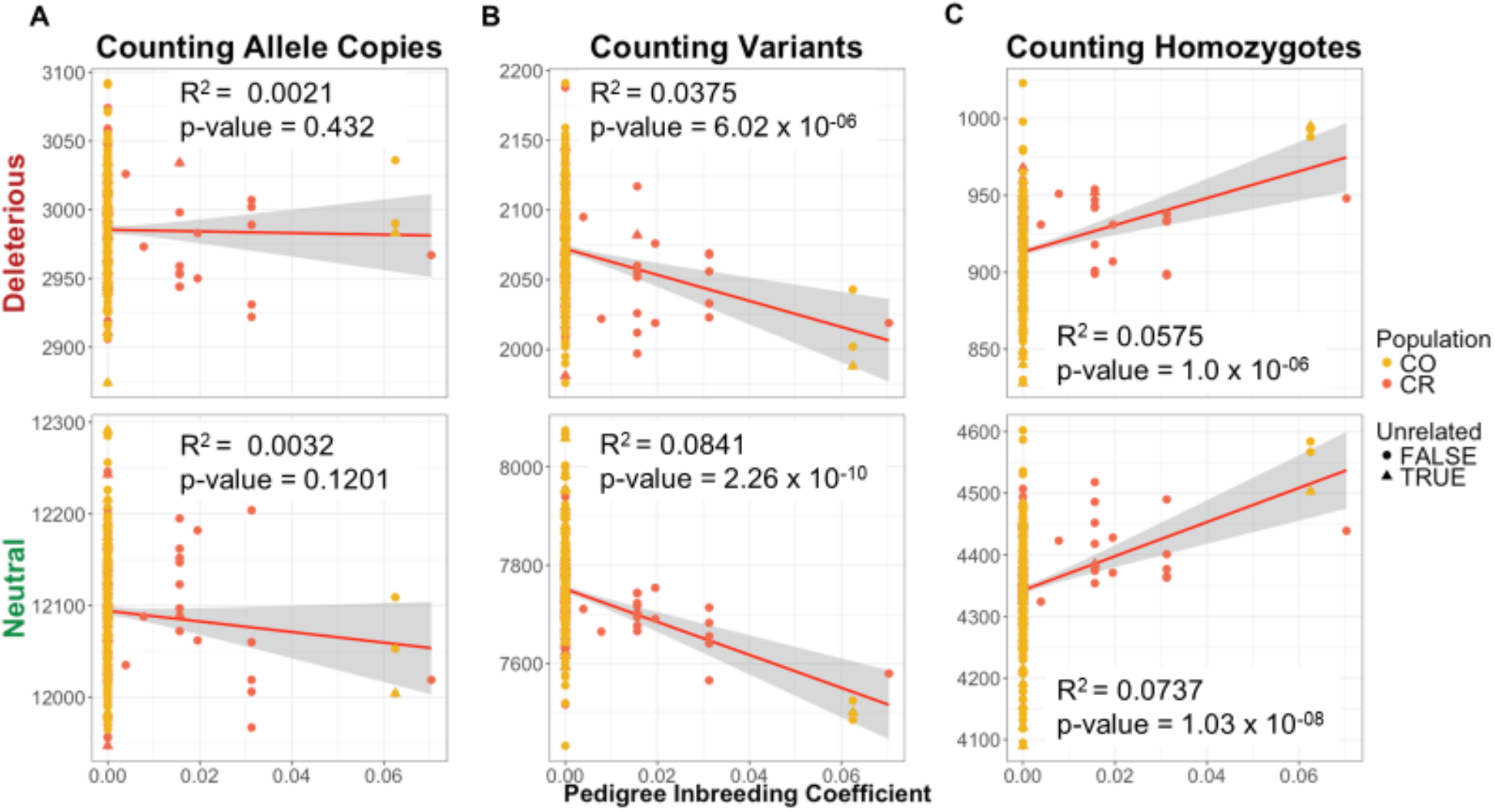
Pedigree inbreeding coefficient (F_PED_) is correlated with deleterious variation. Triangles represent the individuals that were sampled in the unrelated data set (*n*=30). Variants were predicted as either putatively deleterious (nonsynonymous) SNPs or putatively neutral (synonymous) SNPs using GERP^42^. Correlation between F_PED_ and the number of mutations per individual in Colombians and Costa Ricans. (A) Number of derived alleles per individual. (B) Number of variants per individual. (C) Number of homozygous derived genotypes per individual. The first row depicts the correlation between deleterious sites using each counting method and F_PED_ for sequenced individuals from Latin American isolates. The second row depicts the correlation between neutral sites using each counting method and F_PED_ in the same individuals. Population abbreviations are as in **Figure 1**.

## Discussion

Here we present the first comprehensive study of genetic diversity, demographic history, identity-by-descent, runs of homozygosity, and deleterious mutations in multiple admixed isolated populations. We show that admixture sufficiently increases genetic diversity of the Colombian and Costa Rican isolates, such that each isolate has diversity levels comparable to a non-isolated population. However, we still observe characteristics in the Latin American isolates that are hallmarks of an archetypal isolate, such as: an excess of IBD segments, cryptic relatedness within the population, and an enrichment of long ROH. Further, we demonstrate that long ROHs contain an enrichment of deleterious variants carried in the homozygous state, which has potential implications for fitness and disease risk.

Taken together our results support historical data which states that a recent admixture event, within the last 500 years, founded the Colombian and Costa Rican population isolates. A bottleneck corresponding to the Spanish Settlement, followed the founding event and then each population has increased in size until the present day^23,67^. We see evidence of these processes in the inference of demography from IBD patterns. The IBDNe inference shows a severe population bottleneck occurring approximately 500 years ago, coinciding with the historical record^23^. Importantly, the bottleneck experienced in the Latin American isolates was not as prolonged as that experienced by the Finnish. Further, the Finnish bottleneck occurred thousands of years ago. The difference in bottleneck timescales likely accounts for some portion of the higher genetic diversity observed in Latin American population isolates in comparison to the Finnish. In other words, the bottlenecks captured by IBDNe in the Latin Americans are too recent to markedly impact levels of genetic diversity. Further, the admixture process experienced by the Latin American isolates could increase levels of genetic diversity, especially because some individuals have appreciable levels of African ancestry^50^. We see little difference in patterns of genetic variation in the 1000 Genomes Colombian samples and the Colombian sample studied in this project. The Latin American isolates occupy areas that were considered as being geographically isolated at the time of sampling – the Central Valley of Costa Rica and the department of Antioquia in Colombia^23^, while the 1000 Genomes CLM sample was taken from Medellín which is included within the Antioquia region^68–71^. Thus, there is likely some amount of shared demography between the 1000 Genome CLM and our isolated Colombia population. However, it is worth noting that the individuals in the Latin American isolates are from pedigrees ascertained for Bipolar Disorder 1, rather than a random sample from the area.

Our results beg the question, what constitutes a population isolate? For example, is it a requirement that population isolates have low genetic diversity relative to the source population? Under this definition, the Latin American population isolates would not qualify as population isolates. The bottleneck in the Costa Ricans and Colombians seems to have had little effect on their genetic diversity, as their diversity levels are similar to non-isolated Latin American populations. The Finnish, on the other hand, experienced a long-term bottleneck that has resulted in a depletion of segregating sites, and of the remaining segregating sites, there is an enrichment of deleterious variants relative to non-isolated populations^2,13^, and would clearly qualify as an isolate. However, if one measures isolation based on IBD we see that there is an enrichment of IBD segments in the Latin American isolates relative to the Finnish. Further, looking at ROH, Latin American individuals from population isolates have a larger burden of ROH than Finnish, thus increasing the chances of identifying more shared genomic regions in the Latin American isolates than the Finnish. By this metric, the Latin American population isolates would certainly qualify as a population isolate. Thus, both the Costa Rican and Colombian populations and the Finnish are isolates but in different ways. For example, the Costa Ricans and Colombians are historical isolates, meaning these populations are not currently isolated but they exhibit many traits of an isolate, whereas the Finnish are contemporary isolates, meaning the population is still isolated and is the archetypal isolate that one would imagine. Our work also suggests that isolated populations have distinct demographic histories that impact genetic variation in different ways; and it is critical that researchers study and quantify the consequences of demography in each population.

We find that Latin American isolates have the largest ROH burden in comparison to any other sampled population. Our work corroborate results from a recent review on ROH where authors state that populations with small Ne and recent consanguinity will harbor the largest amount of ROH^72^. Because previous research has shown a strong correlation between recent inbreeding, quantified by both F_SNP_ and F_PED_, and long runs of homozygosity, we were particularly interested in the mechanism behind the generation of long ROH^19–21,56–58,73^. We used simulations to test which demographic scenarios could produce long ROH (**Figure 5**). These simulations and availability of extended pedigree data were crucial, because the F_SNP_ metric can also be influenced by a recent bottleneck. Thus, having F_PED_ available allowed us to test whether the correlation we observed between F_SNP_ and ROH was a consequence of a recent bottleneck or recent consanguinity. If small population size or admixture was responsible for generating the ROHs, these processes would not be reflected in F_PED_. Thus, we would not expect to find a correlation between F_PED_ and the amount of the genome in ROHs. The fact that we observe a correlation between F_PED_ and the amount of the genome in ROH suggests that recent consanguinity (as measured by F_PED_) is related to the extent of long ROHs in the genome. Further, our simulations show that neither admixture nor a recent population bottleneck could generate the high levels of long ROH that are observed in some individuals. It was only when we incorporated inbreeding into the simulation that levels of ROH comparable to what we observed in our data were produced. Thus, both lines of evidence suggest that the Latin American population isolates have experienced more recent consanguinity than other population isolates, like the Finnish. Further, in Finland it has previously been shown that the frequency of consanguinity, due to first-cousin marriages, is quite low and the best predictors of these unions were socio-economic class and ethnicity, rather than geographic barriers or population density^74^. On the other hand, for the two Latin American isolates consanguinity could be a consequence of increased geographic barriers preventing movement of individuals over more dispersed areas. It is also important to point out that it is unclear the extent to which ascertaining individuals from large pedigrees may impact the number of ROHs in our sample. Thus, the finding of an increase in ROHs may not be generalizable to Colombian and Costa Rican populations as a whole. However, we observed a similar pattern of increased ROH in the CLM, which suggests that the pedigree ascertainment of the CO and CR may not be generating the increase in ROHs.

We also tested how recent consanguinity affects deleterious variation in the genome. When counting homozygous derived deleterious genotypes, we found a positive correlation between the number of nonsynonymous homozygous genotypes and the amount of an individual’s genome within an ROH (**Figure 6**). Further, we observed an enrichment for nonsynonymous homozygous derived genotypes relative to synonymous homozygous derived genotypes within ROHs versus the rest of the genome (**Table 1**). This enrichment can be a result of nonsynonymous mutations generally segregating at lower frequency and typically being carried as a single copy in an individual. When an ROH is formed, the chromosome that was carrying the mutation is copied, thus allowing the mutation to increase the number of homozygotes within the ROH^19,20^. Since long ROH are a product of recent consanguinity, and these populations have experienced recent consanguinity, we see a corresponding increase in the burden of deleterious variants in the genomes of Costa Rican and Colombian isolates. Since we are more likely to see deleterious variants in the homozygous form in areas of the genome that fall within an ROH, our work is particularly relevant for alleles associated with recessive diseases. Lastly, we provide a mechanism for how recent consanguinity can reduce fitness in natural populations^75–77^ Specifically, if gene-knockouts and deleterious mutations tend to be recessive^40,78–82^, as suggested by several studies, then recent consanguinity will increase the number of homozygous derived deleterious variants carried by an individual in a long ROH, thus leading to a reduction of fitness in the sampled population^6^.

Utilizing estimated ancestry proportions from across the genome, we tested for a correlation between an individual’s ancestry and the amount of their genome that falls within an ROH. To our knowledge, this is the first time that the relationship between proportion of ancestry and the amount of the genome within an ROH has been examined (**Supplementary Figure 8**). We found a positive correlation between the proportion of European ancestry and the amount of an individual’s genome within a run of homozygosity. These results are consistent with the Latin American isolates originating from a small number of European founders, which would decrease genetic diversity and increase homozygosity for those areas of the genome containing European haplotypes. We observed a negative correlation between Native American ancestry and the amount of the genome contained within an ROH (**Supplementary Figure 8**). This finding appears to be at odds with previous research^19,72^ that detected the opposite pattern. Some of this difference may be due to distinct sampling strategies of the Native American source population in our study compared to previous work. The reference Native American population we used was composed of Chibchan-speaking individuals from Reich et al.^48^. Chibchan-speaking populations inherited their Native American ancestry from admixture between Southern and Northern American lineages, the necessity of admixture was particularly apparent in the Cabecar of Costa Rica^48^. Because our reference Native American population is admixed, and Native American populations tend to be small, it is likely that drift has affected different alleles in source populations that formed the current Chibchan-speaking populations. The Chibchan speaking populations may have more diversity, fewer fixed homozygous sites, than previously sampled Native American populations which could explain the negative correlation we observed between ancestry and ROH.

While we were able to capture evidence of recent bottlenecks and expansions within Latin American isolates using IBDNe^35^ (**Figure 3**), our demographic inferences have some limitations. For example, the current estimates of N_e_ are unrealistically large. This inflation may be due to low sample size, since we only used 30 individuals, or it may be a result of applying IBDNe to admixed populations. IBDNe was designed to be applied to un-admixed, randomly mating populations. Thus, one area of future research could be exploring the influence of admixture on IBD and developing methods to infer demography using IBD patterns in admixed populations. Interestingly, in our study, the populations with the highest IBD scores were admixed (PUR, CO, CR, and CLM). Furthermore, because IBD segments may contain useful information for identifying regions of the genome that contain disease associated mutations, especially within individuals with the highest amounts of consanguinity, it may be useful to deconvolute ancestry for each segment when identifying disease associated mutations because disease prevalence may differ in each parental population.

Population isolates have frequently been used for mapping Mendelian disease genes^17,83–88^ and studying complex diseases^16,89–95^. Isolates are thought to be beneficial in comparison to non-isolated populations because of their increased homogeneity of the gene pool, disease causing alleles potentially existing at an appreciable frequency due to drift, possible enrichment in prevalence of the phenotype of interest^1–4^, and a likely reduction in the variability of phenotypes. Our work shows that the genetic diversity and genomic background of population isolates varies immensely. Therefore, it is imperative that we understand the unique genetic diversity belonging to each population isolate. Researchers should adapt their study design to integrate the demographic history of the population, to better leverage the power of the unique genetic features of the population of interest. For example, if we knew beforehand that there was a history of consanguineous unions within the study population, researchers could target ROH for disease mapping. This method has previously been used to identify human knockouts, discover novel loci associated with disease, and understand gene function^95–98^. Further, populations with large amounts of ROH could help us better understand disease architecture because ROH may harbor more recessive mutations that do not have full penetrance, since the prevalence of recessively acting variants in ROH is enriched relative to non-ROH portions of the genome. Most importantly, our work highlights the importance of understanding the demographic history of isolated populations, as differences in demographic history will greatly impact patterns of genetic variation in isolates.

## Supplemental Data

Supplemental data include seventeen figures and two tables.

**Conflicts of Interest:**

## Acknowledgements

The authors would like to acknowledge Charleston Chiang, Jesse Garcia, Malika Kumar, and Sonya McKeown for contributing their time and thoughtful discussion. This material is based upon work supported by the National Science Foundation Graduate Research Fellowship under Grant Numbers DGE-1144087 and DGE-1650604 awarded to JAM, as well as partial support from NIH grant MH095454 awarded to NF, and NIH grant R35 GM119856 awarded to KEL.

## Web Resources

6-primate EPO alignment: ftp://ftp.ensembl.org/pub/release-75/fasta/ancestral_alleles/

ADMIXTURE: https://www.genetics.ucla.edu/software/admixture/download.html

IBDSeq: http://faculty.washington.edu/browning/ibdseq.html

IBDNe: http://faculty.washington.edu/browning/ibdne.html#download

KING (version 2.1): http://people.virginia.edu/~wc9c/KING/history.htm

LAMP: http://lamp.icsi.berkeley.edu/lamp/

PLINK: http://www.cog-genomics.org/plink2

ROH simulation script: https://github.com/LohmuellerLab/ROH_Latin_American_Isolates

SeattleSeq Annotation website: http://snp.gs.washington.edu/SeattleSeqAnnotation138/

SLIM: https://messerlab.org/slim/

VCFTools: http://vcftools.sourceforge.net/downloads.html

GATK: https://software.broadinstitute.org/gatk/download/archive

## References

1. Peltonen, L., Palotie, A., and Lange, K. (2000). Use of population isolates for mapping complex traits. Nat. Rev. Genet. 1, 182–190.

2. Lim, E.T., Würtz, P., Havulinna, A.S., Palta, P., Tukiainen, T., Rehnström, K., Esko, T., Mägi, R., Inouye, M., Lappalainen, T., et al. (2014). Distribution and medical impact of loss-of-function variants in the Finnish founder population. PLoS Genet 10, e1004494.

3. Gudbjartsson, D.F., Helgason, H., Gudjonsson, S.A., Zink, F., Oddson, A., Gylfason, A., Besenbacher, S., Magnusson, G., Halldorsson, B.V., and Hjartarson, E. (2015). Large-scale whole-genome sequencing of the Icelandic population. Nat. Genet. 47, 435.

4. Scott, E.M., Halees, A., Itan, Y., Spencer, E.G., He, Y., Azab, M.A., Gabriel, S.B., Belkadi, A., Boisson, B., Abel, L., et al. (2016). Characterization of Greater Middle Eastern genetic variation for enhanced disease gene discovery. Nat. Genet.

5. Lohmueller, K.E., Indap, A.R., Schmidt, S., Boyko, A.R., Hernandez, R.D., Hubisz, M.J., Sninsky, J.J., White, T.J., Sunyaev, S.R., and Nielsen, R. (2008). Proportionally more deleterious genetic variation in European than in African populations. Nature 451, 994.

6. Charlesworth, D., and Willis, J.H. (2009). The genetics of inbreeding depression. Nat. Rev. Genet. 10, 783.

7. Lohmueller, K.E. (2014). The Impact of Population Demography and Selection on the Genetic Architecture of Complex Traits. PLOS Genet. 10, e1004379.

8. Simons, Y.B., Turchin, M.C., Pritchard, J.K., and Sella, G. (2014). The deleterious mutation load is insensitive to recent population history. Nat. Genet. 46, 220–224.

9. Service, S., DeYoung, J., Karayiorgou, M., Roos, J.L., Pretorious, H., Bedoya, G., Ospina, J., Ruiz-Linares, A., Macedo, A., Palha, J.A., et al. (2006). Magnitude and distribution of linkage disequilibrium in population isolates and implications for genome-wide association studies. Nat. Genet. 38, 556–560.

10. Xue, Y., Mezzavilla, M., Haber, M., McCarthy, S., Chen, Y., Narasimhan, V., Gilly, A., Ayub, Q., Colonna, V., Southam, L., et al. (2017). Enrichment of low-frequency functional variants revealed by whole-genome sequencing of multiple isolated European populations. Nat. Commun. 8, 15927.

11. Kittles, R.A., Perola, M., Peltonen, L., Bergen, A.W., Aragon, R.A., Virkkunen, M., Linnoila, M., Goldman, D., and Long, J.C. (1998). Dual origins of Finns revealed by Y chromosome haplotype variation. Am. J. Hum. Genet. 62, 1171–1179.

12. Peltonen, L., Jalanko, A., and Varilo, T. (1999). Molecular genetics the Finnish disease heritage. Hum. Mol. Genet. 8, 1913–1923.

13. Wang, S.R., Agarwala, V., Flannick, J., Chiang, C.W.K., Altshuler, D., Flannick, J., Manning, A., Hartl, C., Agarwala, V., Fontanillas, P., et al. (2014). Simulation of Finnish Population History, Guided by Empirical Genetic Data, to Assess Power of Rare-Variant Tests in Finland. Am. J. Hum. Genet. 94, 710–720.

14. De La Chapelle, A., and Wright, F.A. (1998). Linkage disequilibrium mapping in isolated populations: the example of Finland revisited. Proc. Natl. Acad. Sci. 95, 12416–12423.

15. Martin, A.R., Karczewski, K.J., Kerminen, S., Kurki, M., Sarin, A.-P., Artomov, M., Eriksson, J.G., Esko, T., Genovese, G., Havulinna, A.S., et al. (2017). Haplotype sharing provides insights into fine-scale population history and disease in Finland. BioRxiv 200113.

16. Panoutsopoulou, K., Hatzikotoulas, K., Xifara, D.K., Colonna, V., Farmaki, A.-E., Ritchie, G.R.S., Southam, L., Gilly, A., Tachmazidou, I., Fatumo, S., et al. (2014). Genetic characterization of Greek population isolates reveals strong genetic drift at missense and trait-associated variants. Nat. Commun. 5, 5345.

17. Nakatsuka, N., Moorjani, P., Rai, N., Sarkar, B., Tandon, A., Patterson, N., Bhavani, G.S., Girisha, K.M., Mustak, M.S., and Srinivasan, S. (2017). The promise of discovering population-specific disease-associated genes in South Asia. Nat. Genet. 49, 1403.

18. Pedersen, C.-E.T., Lohmueller, K.E., Grarup, N., Bjerregaard, P., Hansen, T., Siegismund, H. R., Moltke, I., and Albrechtsen, A. (2017). The Effect of an Extreme and Prolonged Population Bottleneck on Patterns of Deleterious Variation: Insights from the Greenlandic Inuit. Genetics 205, 787–801.

19. Pemberton, T.J., Absher, D., Feldman, M.W., Myers, R.M., Rosenberg, N.A., and Li, J.Z. (2012). Genomic patterns of homozygosity in worldwide human populations. Am. J. Hum. Genet. 91, 275–292.

20. Pemberton, T.J., and Szpiech, Z.A. (2018). Relationship between Deleterious Variation, Genomic Autozygosity, and Disease Risk: Insights from The 1000 Genomes Project. Am. J. Hum. Genet. 0,.

21. Kang, J.T., Goldberg, A., Edge, M.D., Behar, D.M., and Rosenberg, N.A. (2016). Consanguinity Rates Predict Long Runs of Homozygosity in Jewish Populations. Hum. Hered. 82, 87–102.

22. Gaetano, C.D., Fiorito, G., Ortu, M.F., Rosa, F., Guarrera, S., Pardini, B., Cusi, D., Frau, F., Barlassina, C., Troffa, C., et al. (2014). Sardinians Genetic Background Explained by Runs of Homozygosity and Genomic Regions under Positive Selection. PLOS ONE 9, e91237.

23. Carvajal-Carmona, L.G., Ophoff, R., Hartiala, J., Molina, J., Leon, P., Ospina, J., Bedoya, G., Freimer, N., and Ruiz-Linares, A. (2003). Genetic demography of Antioquia (Colombia) and the central valley of Costa Rica. Hum. Genet. 112, 534–541.

24. Consortium, 1000 Genomes Project (2015). A global reference for human genetic variation.

25. Fears, S.C., Kremeyer, B., Araya, C., Araya, X., Bejarano, J., Ramirez, M., Castrillón, G., Gomez-Franco, J., Lopez, M.C., and Montoya, G. (2014). Multisystem component phenotypes of bipolar disorder for genetic investigations of extended pedigrees. JAMA Psychiatry 71, 375–387.

26. Manichaikul, A., Mychaleckyj, J.C., Rich, S.S., Daly, K., Sale, M., and Chen, W.-M. (2010). Robust relationship inference in genome-wide association studies. Bioinformatics 26, 2867–2873.

27. DePristo, M.A., Banks, E., Poplin, R., Garimella, K.V., Maguire, J.R., Hartl, C., Philippakis, A.A., Del Angel, G., Rivas, M.A., and Hanna, M. (2011). A framework for variation discovery and genotyping using next-generation DNA sequencing data. Nat. Genet. 43, 491.

28. Watterson, G.A. (1975). On the number of segregating sites in genetical models without recombination. Theor. Popul. Biol. 7, 256–276.

29. Danecek, P., Auton, A., Abecasis, G., Albers, C.A., Banks, E., DePristo, M.A., Handsaker, R.E., Lunter, G., Marth, G.T., and Sherry, S.T. (2011). The variant call format and VCFtools. Bioinformatics 27, 2156–2158.

30. Browning, B.L., and Browning, S.R. (2013). Detecting Identity by Descent and Estimating Genotype Error Rates in Sequence Data. Am. J. Hum. Genet. 93, 840–851.

31. Browning, S.R., and Browning, B.L. (2007). Rapid and accurate haplotype phasing and missing-data inference for whole-genome association studies by use of localized haplotype clustering. Am. J. Hum. Genet. 81, 1084–1097.

32. Gusev, A., Lowe, J.K., Stoffel, M., Daly, M.J., Altshuler, D., Breslow, J.L., Friedman, J.M., and Pe’er, I. (2009). Whole population, genome-wide mapping of hidden relatedness. Genome Res. 19, 318–326.

33. Browning, B.L., and Browning, S.R. (2013). Improving the accuracy and efficiency of identity-by-descent detection in population data. Genetics 194, 459–471.

34. Kong, A., Gudbjartsson, D.F., Sainz, J., Jonsdottir, G.M., Gudjonsson, S.A., Richardsson, B., Sigurdardottir, S., Barnard, J., Hallbeck, B., and Masson, G. (2002). A high-resolution recombination map of the human genome. Nat. Genet. 31, 241.

35. Browning, S.R., and Browning, B.L. (2015). Accurate non-parametric estimation of recent effective population size from segments of identity by descent. Am. J. Hum. Genet. 97, 404–418.

36. Auton, A., Bryc, K., Boyko, A.R., Lohmueller, K.E., Novembre, J., Reynolds, A., Indap, A., Wright, M.H., Degenhardt, J.D., Gutenkunst, R.N., et al. (2009). Global distribution of genomic diversity underscores rich complex history of continental human populations. Genome Res. 19, 795–803.

37. Sinnwell, J.P., Therneau, T.M., and Schaid, D.J. (2014). The kinship2 R package for pedigree data. Hum. Hered. 78, 91–93.

38. Haller, B.C., and Messer, P.W. (2016). SLiM 2: flexible, interactive forward genetic simulations. Mol. Biol. Evol. 34, 230–240.

39. Kim, B.Y., Huber, C.D., and Lohmueller, K.E. (2017). Inference of the distribution of selection coefficients for new nonsynonymous mutations using large samples. Genetics 206, 345–361.

40. Huber, C.D., Kim, B.Y., Marsden, C.D., and Lohmueller, K.E. (2017). Determining the factors driving selective effects of new nonsynonymous mutations. Proc. Natl. Acad. Sci. 114, 4465–4470.

41. Long, M., and Deutsch, M. (1999). Association of intron phases with conservation at splice site sequences and evolution of spliceosomal introns. Mol. Biol. Evol. 16, 1528–1534.

42. Cooper, G.M., Stone, E.A., Asimenos, G., Green, E.D., Batzoglou, S., and Sidow, A. (2005). Distribution and intensity of constraint in mammalian genomic sequence. Genome Res. 15, 901–913.

43. Alexander, D.H., Novembre, J., and Lange, K. (2009). Fast model-based estimation of ancestry in unrelated individuals. Genome Res. 19, 1655–1664.

44. Sankararaman, S., Sridhar, S., Kimmel, G., and Halperin, E. (2008). Estimating local ancestry in admixed populations. Am. J. Hum. Genet. 82, 290–303.

45. Pagani, L., Clair, P.A.S., Teshiba, T.M., Fears, S.C., Araya, C., Araya, X., Bejarano, J., Ramirez, M., Castrillón, G., and Gomez-Makhinson, J. (2016). Genetic contributions to circadian activity rhythm and sleep pattern phenotypes in pedigrees segregating for severe bipolar disorder. Proc. Natl. Acad. Sci. 113, E754–E761.

46. Consortium, I.H. (2003). The international HapMap project. Nature 426, 789.

47. Consortium, I.H. (2007). A second generation human haplotype map of over 3.1 million SNPs. Nature 449, 851.

48. Reich, D., Patterson, N., Campbell, D., Tandon, A., Mazieres, S., Ray, N., Parra, M.V., Rojas, W., Duque, C., and Mesa, N. (2012). Reconstructing native American population history. Nature 488, 370.

49. Aulchenko, Y.S., Ripke, S., Isaacs, A., and Van Duijn, C.M. (2007). GenABEL: an R library for genome-wide association analysis. Bioinformatics 23, 1294–1296.

50. Lohmueller, K.E., Albrechtsen, A., Li, Y., Kim, S.Y., Korneliussen, T., Vinckenbosch, N., Tian, G., Huerta-Sanchez, E., Feder, A.F., and Grarup, N. (2011). Natural selection affects multiple aspects of genetic variation at putatively neutral sites across the human genome. PLoS Genet. 7, e1002326.

51. Cai, J.J., Macpherson, J.M., Sella, G., and Petrov, D.A. (2009). Pervasive hitchhiking at coding and regulatory sites in humans. PLoS Genet. 5, e1000336.

52. Hernandez, R.D., Kelley, J.L., Elyashiv, E., Melton, S.C., Auton, A., McVean, G., Sella, G., and Przeworski, M. (2011). Classic selective sweeps were rare in recent human evolution. Science 331, 920–924.

53. Stumpf, M.P., and Goldstein, D.B. (2003). Demography, recombination hotspot intensity, and the block structure of linkage disequilibrium. Curr. Biol. 13, 1–8.

54. Pritchard, J.K., and Przeworski, M. (2001). Linkage disequilibrium in humans: models and data. Am. J. Hum. Genet. 69, 1–14.

55. Gravel, S., Zakharia, F., Moreno-Estrada, A., Byrnes, J.K., Muzzio, M., Rodriguez-Flores, J.L., Kenny, E.E., Gignoux, C.R., Maples, B.K., Guiblet, W., et al. (2013). Reconstructing Native American Migrations from Whole-Genome and Whole-Exome Data. PLOS Genet. 9, e1004023.

56. Szpiech, Z.A., Xu, J., Pemberton, T.J., Peng, W., Zöllner, S., Rosenberg, N.A., and Li, J.Z. (2013). Long runs of homozygosity are enriched for deleterious variation. Am. J. Hum. Genet. 93, 90–102.

57. McQuillan, R., Leutenegger, A.-L., Abdel-Rahman, R., Franklin, C.S., Pericic, M., Barac-Lauc, L., Smolej-Narancic, N., Janicijevic, B., Polasek, O., and Tenesa, A. (2008). Runs of homozygosity in European populations. Am. J. Hum. Genet. 83, 359–372.

58. Kirin, M., McQuillan, R., Franklin, C.S., Campbell, H., McKeigue, P.M., and Wilson, J.F. (2010). Genomic runs of homozygosity record population history and consanguinity. PloS One 5, e13996.

59. Pemberton, T.J., and Szpiech, Z.A. (2018). Relationship between Deleterious Variation, Genomic Autozygosity, and Disease Risk: Insights from The 1000 Genomes Project. Am. J. Hum. Genet. 0,.

60. Ceballos, F.C., Joshi, P.K., Clark, D.W., Ramsay, M., and Wilson, J.F. (2018). Runs of homozygosity: windows into population history and trait architecture. Nat. Rev. Genet.

61. Kardos, M., Taylor, H.R., Ellegren, H., Luikart, G., and Allendorf, F.W. (2016). Genomics advances the study of inbreeding depression in the wild. Evol. Appl. 9, 1205–1218.

62. Kimura, M., Maruyama, T., and Crow, J.F. (1963). The mutation load in small populations. Genetics 48, 1303–1312.

63. Ohta, T. (1973). Slightly deleterious mutant substitutions in evolution. Nature 246, 96.

64. Hodgkinson, A., Casals, F., Idaghdour, Y., Grenier, J.-C., Hernandez, R.D., and Awadalla, P. (2013). Selective constraint, background selection, and mutation accumulation variability within and between human populations. BMC Genomics 14, 495.

65. Peischl, S., Dupanloup, I., Kirkpatrick, M., and Excoffier, L. (2013). On the accumulation of deleterious mutations during range expansions. Mol. Ecol. 22, 5972–5982.

66. Fu, W., Gittelman, R.M., Bamshad, M.J., and Akey, J.M. (2014). Characteristics of Neutral and Deleterious Protein-Coding Variation among Individuals and Populations. Am. J. Hum. Genet. 95, 421–436.

67. Escamilla, M.A., Spesny, M., Reus, V.I., Gallegos, A., Meza, L., Molina, J., Sandkuijl, L.A., Fournier, E., Leon, P.E., Smith, L.B., et al. (1996). Use of linkage disequilibrium approaches to map genes for bipolar disorder in the Costa Rican population. Am. J. Med. Genet. 67, 244–253.

68. Wang, S., Ray, N., Rojas, W., Parra, M.V., Bedoya, G., Gallo, C., Poletti, G., Mazzotti, G., Hill, K., and Hurtado, A.M. (2008). Geographic patterns of genome admixture in Latin American Mestizos. PLoS Genet. 4, e1000037.

69. Bedoya, G., Montoya, P., García, J., Soto, I., Bourgeois, S., Carvajal, L., Labuda, D., Alvarez, V., Ospina, J., and Hedrick, P.W. (2006). Admixture dynamics in Hispanics: a shift in the nuclear genetic ancestry of a South American population isolate. Proc. Natl. Acad. Sci. 103, 7234–7239.

70. Safford, F., and Palacios, M. (2002). Colombia: Fragmented land, divided society (Oxford University Press, USA).

71. Carvajal-Carmona, L.G., Soto, I.D., Pineda, N., Ortíz-Barrientos, D., Duque, C., Ospina-Duque, J., McCarthy, M., Montoya, P., Alvarez, V.M., and Bedoya, G. (2000). Strong Amerind/white sex bias and a possible Sephardic contribution among the founders of a population in northwest Colombia. Am. J. Hum. Genet. 67, 1287–1295.

72. Ceballos, F.C., Joshi, P.K., Clark, D.W., Ramsay, M., and Wilson, J.F. (2018). Runs of homozygosity: windows into population history and trait architecture. Nat. Rev. Genet.

73. Li, L.-H., Ho, S.-F., Chen, C.-H., Wei, C.-Y., Wong, W.-C., Li, L.-Y., Hung, S.-I., Chung, W.-H., Pan, W.-H., Lee, M.-T.M., et al. (2006). Long contiguous stretches of homozygosity in the human genome. Hum. Mutat. 27, 1115–1121.

74. Jorde, L.B., and Pitkänen, K.J. (1991). Inbreeding in Finland. Am. J. Phys. Anthropol. 84, 127–139.

75. Wright, S. (1984). Evolution and the genetics of populations, volume 3: experimental results and evolutionary deductions (University of Chicago press).

76. Charlesworth, B., and Charlesworth, D. (1999). The genetic basis of inbreeding depression. Genet. Res. 74, 329–340.

77. Wang, J., Hill, W.G., Charlesworth, D., and Charlesworth, B. (1999). Dynamics of inbreeding depression due to deleterious mutations in small populations: mutation parameters and inbreeding rate. Genet. Res. 74, 165–178.

78. Balick, D.J., Do, R., Cassa, C.A., Reich, D., and Sunyaev, S.R. (2015). Dominance of deleterious alleles controls the response to a population bottleneck. PLoS Genet. 11, e1005436.

79. Mukai, T., Chigusa, S.I., Mettler, L.E., and Crow, J.F. (1972). Mutation rate and dominance of genes affecting viability in Drosophila melanogaster. Genetics 72, 335–355.

80. Simmons, M.J., and Crow, J.F. (1977). Mutations affecting fitness in Drosophila populations. Annu. Rev. Genet. 11, 49–78.

81. Phadnis, N., and Fry, J.D. (2005). Widespread correlations between dominance and homozygous effects of mutations: implications for theories of dominance. Genetics 171, 385–392.

82. Agrawal, A.F., and Whitlock, M.C. (2011). Inferences about the distribution of dominance drawn from yeast gene knockout data. Genetics 187, 553–566.

83. Myerowitz, R., and Costigan, F.C. (1988). The major defect in Ashkenazi Jews with Tay-Sachs disease is an insertion in the gene for the alpha-chain of beta-hexosaminidase. J. Biol. Chem. 263, 18587–18589.

84. Hästbacka, J., de la Chapelle, A., Mahtani, M.M., Clines, G., Reeve-Daly, M.P., Daly, M., Hamilton, B.A., Kusumi, K., Trivedi, B., and Weaver, A. (1994). The diastrophic dysplasia gene encodes a novel sulfate transporter: positional cloning by fine-structure linkage disequilibrium mapping. Cell 78, 1073–1087.

85. Aittomäki, K., Lucena, J.D., Pakarinen, P., Sistonen, P., Tapanainen, J., Gromoll, J., Kaskikari, R., Sankila, E.-M., Lehväslaiho, H., and Engel, A.R. (1995). Mutation in the follicle-stimulating hormone receptor gene causes hereditary hypergonadotropic ovarian failure. Cell 82, 959–968.

86. Ruiz-Perez, V.L., Ide, S.E., Strom, T.M., Lorenz, B., Wilson, D., Woods, K., King, L., Francomano, C., Freisinger, P., and Spranger, S. (2000). Mutations in a new gene in Ellis-van Creveld syndrome and Weyers acrodental dysostosis. Nat. Genet. 24, 283.

87. Verhoeven, K., Villanova, M., Rossi, A., Malandrini, A., De Jonghe, P., and Timmerman, V. (2001). Localization of the gene for the intermediate form of Charcot-Marie-Tooth to chromosome 10q24. 1-q25. 1. Am. J. Hum. Genet. 69, 889–894.

88. Valente, E.M., Bentivoglio, A.R., Dixon, P.H., Ferraris, A., Ialongo, T., Frontali, M., Albanese, A., and Wood, N.W. (2001). Localization of a novel locus for autosomal recessive early-onset parkinsonism, PARK6, on human chromosome 1p35-p36. Am. J. Hum. Genet. 68, 895–900.

89. McInnes, L.A., Reus, V.I., Barnes, G., Charlat, O., Jawahar, S., Lewitzky, S., Yang, Q., Duong, Q., Spesny, M., and Araya, C. (2001). Fine-scale mapping of a locus for severe bipolar mood disorder on chromosome 18p11. 3 in the Costa Rican population. Proc. Natl. Acad. Sci. 98, 11485–11490.

90. Ober, C., Tan, Z., Sun, Y., Possick, J.D., Pan, L., Nicolae, R., Radford, S., Parry, R.R., Heinzmann, A., and Deichmann, K.A. (2008). Effect of variation in CHI3L1 on serum YKL-40 level, risk of asthma, and lung function. N. Engl. J. Med. 358, 1682–1691.

91. Stacey, S.N., Manolescu, A., Sulem, P., Thorlacius, S., Gudjonsson, S.A., Jonsson, G.F., Jakobsdottir, M., Bergthorsson, J.T., Gudmundsson, J., Aben, K.K., et al. (2008). Common variants on chromosome 5p12 confer susceptibility to estrogen receptor-positive breast cancer. Nat. Genet. 40, 703–706.

92. Stacey, S.N., Sulem, P., Jonasdottir, A., Masson, G., Gudmundsson, J., Gudbjartsson, D.F., Magnusson, O.T., Gudjonsson, S.A., Sigurgeirsson, B., and Thorisdottir, K. (2011). A germline variant in the TP53 polyadenylation signal confers cancer susceptibility. Nat. Genet. 43, 1098.

93. Gudmundsson, J., Sulem, P., Gudbjartsson, D.F., Masson, G., Agnarsson, B.A., Benediktsdottir, K.R., Sigurdsson, A., Magnusson, O.T., Gudjonsson, S.A., and Magnusdottir, D.N. (2012). A study based on whole-genome sequencing yields a rare variant at 8q24 associated with prostate cancer. Nat. Genet. 44, 1326.

94. Tachmazidou, I., Dedoussis, G., Southam, L., Farmaki, A.-E., Ritchie, G.R., Xifara, D.K., Matchan, A., Hatzikotoulas, K., Rayner, N.W., and Chen, Y. (2013). A rare functional cardioprotective APOC3 variant has risen in frequency in distinct population isolates. Nat. Commun. 4, 2872.

95. Saleheen, D., Natarajan, P., Armean, I.M., Zhao, W., Rasheed, A., Khetarpal, S.A., Won, H-H., Karczewski, K.J., O’Donnell-Luria, A.H., and Samocha, K.E. (2017). Human knockouts and phenotypic analysis in a cohort with a high rate of consanguinity. Nature 544, 235.

96. Botstein, D., and Risch, N. (2003). Discovering genotypes underlying human phenotypes: past successes for mendelian disease, future approaches for complex disease. Nat. Genet. 33, 228.

97. Lencz, T., Lambert, C., DeRosse, P., Burdick, K.E., Morgan, T.V., Kane, J.M., Kucherlapati, R., and Malhotra, A.K. (2007). Runs of homozygosity reveal highly penetrant recessive loci in schizophrenia. Proc. Natl. Acad. Sci. 104, 19942–19947.

98. Mezzavilla, M., Vozzi, D., Badii, R., Alkowari, M.K., Abdulhadi, K., Girotto, G., and Gasparini, P. (2015). Increased rate of deleterious variants in long runs of homozygosity of an inbred population from Qatar. Hum. Hered. 79, 14–19.

